# A post-mortem investigation of the locus coeruleus-noradrenergic system in resilience to childhood abuse

**DOI:** 10.1101/2025.03.22.644719

**Authors:** Déa Slavova, Maria Antonietta Davoli, Celine Keime, Gabriella Frosi, Erika Vigneault, Corina Nagy, Gustavo Turecki, Bruno Giros, Naguib Mechawar, Elsa Isingrini

## Abstract

Childhood abuse (CA) is one of the strongest lifetime predictors of major depressive disorder (MDD) and suicide. However, some individuals exposed to CA are resilient, avoiding the development of psychopathology. Recently, the locus coeruleus-noradrenergic (LC-NE) system has been involved in resilience following stressful events at adulthood. We investigated how a history of CA affects the integrity of the LC-NE system at the molecular and cellular level in human post-mortem brain samples of depressed suicides, and whether differential neurobiological mechanisms can be revealed in resilient individuals. Anatomical analysis revealed that CA-induced MDD and suicide is associated with decrease in LC-NE neurons density. RNA sequencing of laser captured LC-NE neurons highlighted differentially expressed genes, principally in the RES-CA group. Resilience to CA involves specific neurobiological adaptations in the LC-NE system that potentially protect against the loss of LC-NE neurons and the negative long-term outcome of CA-induced depression and suicide. Our results provide insights into potential therapeutic targets for preventing or treating CA-induced MDD.

## Introduction

Major depressive disorder (MDD), associated with loss of motivation and anhedonia, (1), is the leading cause of disability worldwide. MDD is currently one of the strongest predictors of suicide (2), and its development is enforced by a history of child abuse (CA). CA, defined as emotional, physical and/or sexual maltreatment or abuse has devastating and lasting impact on psychology (3), physiology (4) and brain structure, function and connectivity (5). Clinical observations suggest that MDD individuals with a history of CA are younger at onset, exhibit greater symptom severity, have poorer treatment response, and show increased risk of suicidal behaviour compared to individuals with MDD without a history of CA (6,7).

However, the long-term consequences of CA vary from one individual to another. While CA can precipitate the emergence of MDD, some individuals escape this condition. Indeed, some children have the ability to adapt in response to significant source of adversity and avoid its negative consequences. This ability is defined as resilience (8). Operationalizing the concept of resilience in the context of CA deserves attention: advances in our understanding of resilience indicate that it is an active and adaptive process, and not simply the absence of a pathological response that occurs in more susceptible individuals (9).

From a neurodevelopmental perspective, CA may contribute to establish dysfunctional neurobiological networks encompassing higher-order regions involved in threat and reward processing, emotion regulation, cognitive and executive control (10). Imaging studies have highlighted the potential effect of CA on the developing brain at the structural, functional and connectivity level (10). These include alterations in the anterior cingulate cortex and extended prefrontal areas, as well as in the limbic system, notably the ventral striatum, hippocampus, and amygdala (5,11). Moreover, CA result in lasting effects on brain systems and circuits that mediate the stress response, including the hypothalamic-pituitary-adrenal (HPA) axis and norepinephrine (NE) systems (5,12,13). This body of evidence supports enduring poor adaptive coping and increased risk of psychiatric disorders.

However, the search for protective factors that yield positive psychiatric outcomes has only recently emerged (14). Imaging studies suggest that resilience to the effects of CA may be mediated by larger hippocampal volume (15), increased fractional anisotropy in the posterior cingulum (16), and positive resting-state functional connectivity between prefrontal areas and limbic regions (17,18). These studies have the potential to advance our understanding of the neural mechanisms underpinning the link between resilience and psychopathology in the context of CA. However, how the resilience network is controlled by stress-integrative structures and the upstream neuronal and molecular mechanisms responsible for the functional alteration of this network remains to be deeply investigated.

The noradrenergic (NE) system comprises clusters of neurons, with the locus coeruleus (LC) being the largest nucleus (19).

NE is released throughout the brain, and its regulation of cortical and limbic areas participates in a variety of behavioral and physiological processes, including arousal and stress reactivity, cognitive and executive function, mood, and emotions. The involvement of the LC-NE system in CA-induced psychopathology could result from altered LC-NE modulation of cognitive and limbic brain areas, consequently inducing cognitive and emotional dysfunction associated with depression and suicide (20–22).

Furthermore, the LC-NE system also plays a critical role in the regulation of brain development (23). It is known to be involved in shaping brain wiring during critical periods of development, where changes in NE neurotransmission can have devastating effects on the development of neuronal networks and permanently alter cognitive and emotional performance. For example, changes in the expression of key genes involved in the regulation of NE neurotransmission in rats during vulnerable developmental periods can cause a range of abnormal behaviors later in life (24). Early stress has also been identified as a disruptor of the LC-NE system at both the anatomical and functional levels (25–27). Early stress is associated with lasting increases in noradrenergic responsivity in preclinical models. Changes in LC-NE neurotransmission associated with early life adversity (ELA) may thus have lasting impacts on cognitive and emotional functions throughout the lifespan, potentially sustaining psychopathological development.

Moreover, in recent years, the LC-NE system has been shown to play an important role in resilience to stress in adulthood in both humans (28,29) and rodents (30,31). However, the implication of the LC-NE system in resilient individuals with a history of CA has not been established, despite the significant effects that CA can have on this system.

In this study, we examined how a history of CA may affect the integrity of the LC-NE system at the molecular and cellular levels in human post-mortem brain samples of depressed suicides, and whether differential neurobiological mechanisms can be revealed in resilient individuals exposed to CA who did not develop any psychopathology. First, we aimed to characterize the impact of CA on the density of NE neurons in the LC using immunostaining. Although recent studies provide important information for understanding the spatial distribution of the LC neurons based on their molecular identity (32,33), the molecular profile of LC-NE cells involved in resilience against CA are still unknown. Consequently, we performed RNA sequencing of isolated NE neurons using laser capture microdissection (LCM) in the LC to identify potential transcriptomic adaptations occurring in LC-NE cells of resilient individuals with a history of CA.

## METHODS

### Human post-mortem samples

Human LC tissues were obtained from the Douglas-Bell Canada Brain Bank (DBCBB, Montreal, Canada). All brains were donated to the Suicide section of the DBCBB by familial consent through the Quebec Coroner’s Office. The left hemisphere was cut into thick (2.5 cm) sections, snap-frozen, and stored at -80°C, while the right hemisphere was fixed by immersion in formalin. Expert staff from the brain bank conducted dissections of the LC under the guidance of human brain atlases (34). The entire LC was collected along the rostro-caudal axis of the brainstem (∼Obex + 32 mm to ∼Obex + 20 mm; **Figure 1E**).

**Figure 1.**
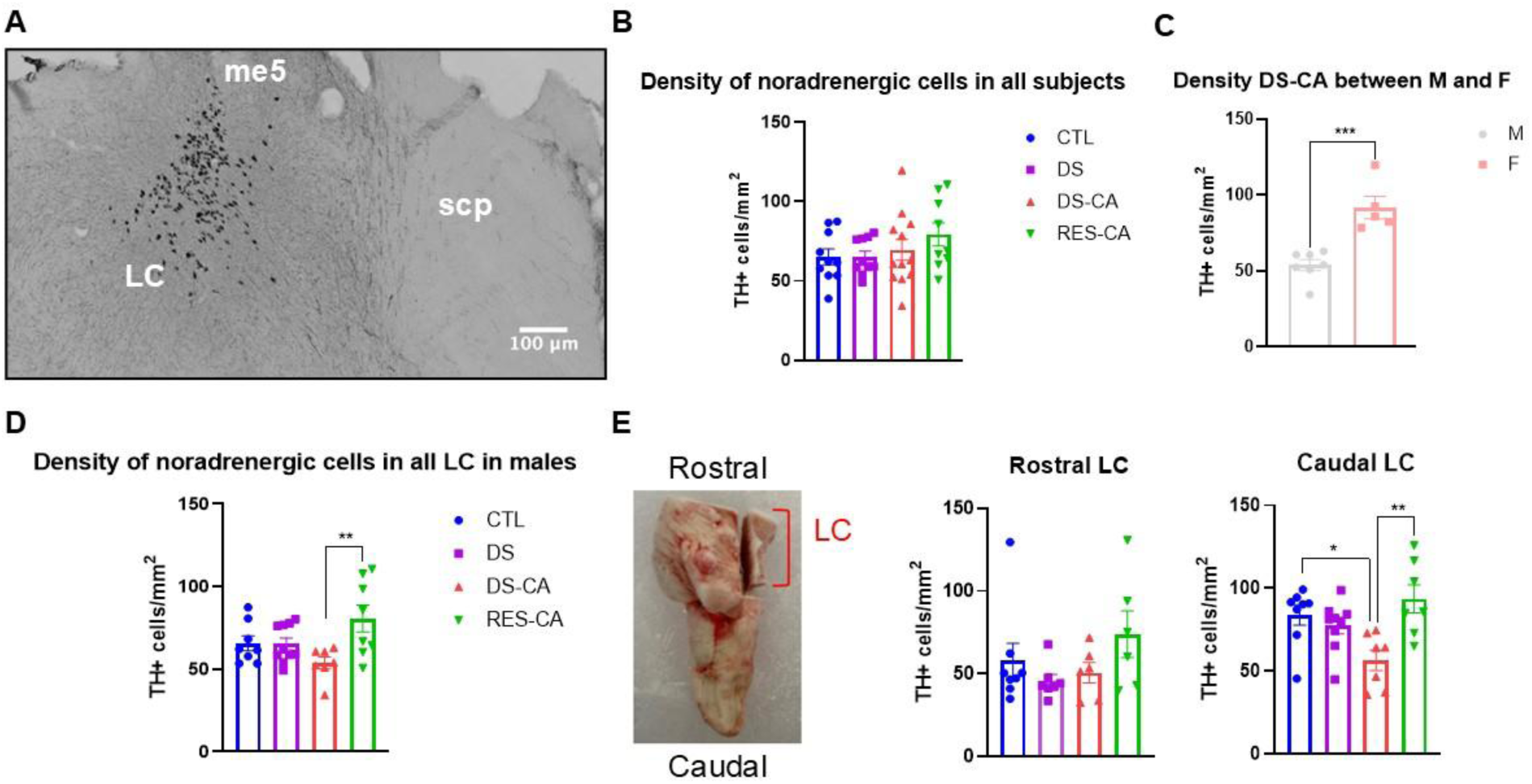
Density of noradrenergic cells is preserved in resilient individuals exposed to child abuse. **A**. Representative photomicrograph of DAB immunostaining against tyrosine hydroxylase (TH), visualizing noradrenergic (NE) neurons in the locus coeruleus (LC) in human post-mortem tissues. The LC is annotated using references from the human brainstem atlas, including the mesencephalic trigeminal tract (me5) and the superior cerebellar peduncle (scp). **B**. Density of TH+ cells in the LC, including male (M) and female (F) subjects, across different groups: Control (CTL, N = 10, with 1 female), Depression-Suicide (DS, N = 10, no females), Depression-Suicide-Child Abuse (DS-CA, N = 12, with 5 females), and Child Abuse-No Depression (RES-CA, N = 9, with 1 female). No significant difference in TH+ cell density is observed between groups (F_3,37_ = 1.17, p = 0.33). **C**. Within the DS-CA group, the density of TH+ cells in the entire LC is significantly higher in females compared to males (t_10_ = 5.065, p = 0.0005). **D**. In males, the density of TH+ cells in the entire LC is significantly lower in the DSCA group compared to the RES-CA group (group effect: F_3,29_ = 4.06, p = 0.016, followed by Tukey’s multiple comparison test). **E**. **Left**: Representative picture of human LC dissection according to the “rostral” and “caudal” orientation; **Middle**: TH+ cell density in the rostral part of the LC in males does not differ significantly between groups (Kruskal-Wallis test: H_3_ = 3.22, p = 0.36, some of the subjects does not have rostral part). **Right**: TH+ cell density in the caudal part of the LC is significantly decreased in the DS-CA group compared to the CTL and RESCA groups (group effect: F_3,27_ = 5.44, p = 0.0047, followed by Tukey’s multiple comparison test). All results are shown as mean +/- SEM. Statistical significance is indicated as follow: adjusted p-value (q) *< 0.05, **< 0.01, ***< 0.001.

Samples used were from suicide completers who died during an episode of MDD with a history of CA (DS-CA) or without (DS), and matched samples from psychiatrically healthy individuals with a history of CA (RES-CA) or without (CTL) who died suddenly without prolonged agonal period, from cardiovascular arrest, or as passengers in car accidents. These four groups were matched according to age, post-mortem interval (PMI), and pH (**Table1**) (35). Phenotypic information was obtained through standardized psychological autopsies as mentioned in (35). This process included DSM-IV Axis I diagnostic information and childhood experience of care and abuse (CECA) interviews (36).

#### Inclusion criteria

*Inclusion criteria for cases and controls:* (1) Absence of organic brain disease; (2) Death not caused by direct brain lesion; (3) Negative evidence of chronic inflammatory illness. *Inclusion criteria for cases:* (1) Current (6-month) diagnosis of major depressive disorder; (2) No current or lifetime history of manic and/or hypomanic episodes; (3) No current or lifetime history of psychotic disorders.

### Immunohistochemistry

Fixed LC samples were cryoprotected in 15% sucrose at 4°C and serially cut in a rostro-caudal orientation into 50µm sections using a microtome. Sections were stored at -20°C in a 30% glycerol - 30% Ethylene glycol - 40%PBS 0.1M solution. A subset of sections at regular intervals (200µm) was processed for immunostaining. A standard antigen retrieval step using Sodium Citrate Buffer (SCB, 10mM, pH 6) was performed followed by endogenous peroxidase inactivation in 3% H_2_O_2_ for 15 minutes. Incubation with a solution of primary antibody (mouse-anti TH, 1:2000, MAB318, Sigma Aldrich) diluted in a blocking solution (BS) of 0.1M PBS containing 2% normal horse serum and 0.3% Triton-X100 was performed overnight at RT under constant agitation. Sections were rinsed and incubated for 1.5 hours at RT under agitation with a secondary antibody solution (Horse Anti-Mouse IgG Antibody (H+L) Biotinylated, 1:500, BA-2000-1.5, Vector Laboratories) diluted in the BS. Sections were rinsed and incubated in an ABC-HRP kit (VECTASTAIN® Elite® PK-6100) for 30 minutes at RT. Labelling was revealed with DAB substrate kit, Peroxidase HRP, with Nickel, (3,3’-diaminobenzidine,SK-4100,maximum 10 minutes). Sections were mounted on Superfrost charged slides and dehydrated for 2 minutes, twice in the following solutions: H_2_Od, 50% Ethanol, 75% Ethanol, 95% Ethanol, 100% Ethanol, Xylene. Finally, sections were cover slipped with hydrophobic mounting medium (Micromount).

Whole LC sections were scanned using a Zeiss Axio Scan 7 with an Axiocam MRM camera at 10x magnification. Sections were annotated based on the Atlas of the Human Brainstem (37), using the mesencephalic trigeminal tract (me5) and superior cerebellar peduncle (scp) as references (**Figure 1A)**. An average of 18 sections per subject were analysed by a blinded researcher using Icy software (https://icy.bioimageanalysis.org/), employing the automatic spot detector and manually verifying the results. Cell density was calculated as the number of TH-positive cells per µm².

### RNA sequencing

#### Laser capture microdissection (LCM)

Frozen samples (**Table1**) were used for laser LCM. 10µm thick coronal sections, spaced 50µm apart from the middle of the LC, were collected on PEN membrane slides (Carl Zeiss) using a cryostat (Leica). Before processing, the slides were allowed to equilibrate at RT for 5 minutes and dehydrated in freshly prepared solutions for 1 minute as follows: ethanol 70%, 95% and, 100%, and xylene for 2 minutes, to minimize RNA degradation. Neuromelanin was used as a natural marker of LC-NE neurons (22). Approximately 300 neurons were collected from each subject into a 0.5 ml microtube with a silicone cap using a Zeiss Palm Microbeam. For details on the procedure and parameters, refer to the **Supplementary Figure 1.**

#### RNA extraction and quality control

Following LCM, 20µl of PicoPure extraction buffer (Thermofisher, KIT0214) was added to the silicone cap containing the extracted neurons. The microtube was then placed in the CapSure–ExtracSure assembly and covered with an incubation block preheated to 42°C for 30 minutes, centrifuged at 800 × g for 2 minutes. RNA extraction was performed using the PicoPure RNA extraction kit (KIT0204). RNA quality was assessed with the High Sensitivity RNA ScreenTape (Agilent, #228267) using the 2200 TapeStation Controller Software. The remaining RNA samples were stored at -80°C until sequencing.

#### Sequencing

Library preparation was performed at the GenomEast platform at the Institute of Genetics and Molecular and Cellular Biology. Full length cDNA was generated from 1ng or less of total RNA using SMART-SeqX v4 UltraX Low Input RNA Kit for Sequencing (Takara Bio Europe, Saint Germain en Laye, France) with 12 cycles of PCR for cDNA amplification by Seq-Amp polymerase. Six hundreds pg of pre-amplified cDNA were then used as input for Tn5 transposon tagmentation by the Nextera XT DNA Library Preparation Kit (Illumina, San Diego, USA) followed by 12 cycles of library amplification. Following purification with SPRIselect beads (Beckman-Coulter, Villepinte, France), the size and concentration of libraries were assessed by capillary electrophoresis. Libraries were sequenced on an Illumina NextSeq 2000 sequencer as single read 50 base reads. Image analysis and base calling were performed using RTA version 2.7.7 and BCL Convert version 3.8.4 (Illumina).

#### Analysis of LCM-sequencing data

Reads were preprocessed using cutadapt (38) version 4.2 to remove adapter, polyA and low-quality sequences (Phred quality score below 20). Reads shorter than 40 bases were discarded. Reads mapping to rRNA were also discarded (bowtie version 2.2.8 (39)). Reads were then mapped onto the GRCh38 human genome assembly using STAR version 2.7.10b (40). Gene expression was quantified using htseq-count version 0.13.5 (41) and gene annotations from Ensembl release 108. Only genes with at least 10 reads in at least 10 samples have been kept for further analysis. Differential gene expression (DEG) analysis was performed using R 4.1.1 and DESeq2 1.34.0 Bioconductor library (42), adjusting for one surrogate variable (SV) estimated using svaseq method from the sva 3.42.0 Bioconductor package (43). To minimize over adjustment bias while preserving biological variability and given the limited number of samples (n=47), only SV1 was used. SV1, accounting for 34% of the variability, was associated with the transcript integrity number (TIN, r = 0.95, p = 0), pH (r = - 0.57, p = 0) and age (r = 0.34, p = 0.02), controlling for the variables most linked to RNA integrity. To ensure the reliability of the DEG results, an additional analysis was conducted by filtering out mitochondrial genes prior to the DEG analysis, revealing highly similar results. Consequently, all genes were included in the final DEG analysis. To visualize the expression profile of a given gene in several groups on heatmaps, z-scores were calculated from read counts normalized with the median of ratio method (implemented in DESeq2 1.34.0 Bioconductor library) and divided by the median of the length in kb of all transcripts corresponding to the gene. All the packages used for gene enrichment analysis were conducted on R-studio.

### Statistical analysis

All statistical analyses were performed using R-Studio and/or Prism 8 (GraphPad Software). For anatomical analysis, the normality of the distribution and homogeneity of variances were tested using the Shapiro-Wilk (QQ plot) and Bartlett’s test (if the normality was strong; otherwise, Leven’s test was used). If these conditions were validated, one-way ANOVA was used, followed by Tukey’s post hoc test to identify the differences between the groups. Otherwise, non-parametric tests were used, such as the Kruskal Wallis test, followed by Dunn’s multiple comparison test. For comparisons between two samples, two-sided Student’s t-test (Test T) was used when the conditions were satisfied. All results are expressed as mean ± SEM (standard error of the mean), with a threshold set of 0.05. For statistical analysis of RNA-seq data, either Wald (for comparisons between two groups) or Likelihood Ratio Test (LRT) (for the comparisons across all groups) from DESeq2 1.34.0 Bioconductor library with Benjamini and Hochberg p-value adjustment method was used (42).

## RESULTS

### Locus coeruleus anatomical features in resilience

To identify anatomical features of the LC-NE system in the context of CA-induced depression or resilience, we performed a quantitative analysis of LC-NE neurons density in the LC. The number of TH+ cells as well as the area of each layer was measured, allowing to generate TH density values (**Figure 1A**).

When including all subjects (male and female), there were no significant differences in the density of NE cells between groups (**Figure 1B**; F_3,37_ = 1.12, p = 0.33). Due to the limited number of female samples in the CTL, DS, and RES-CA groups, a sex-dependent analysis was not feasible. However, within the DS-CA group, a sex comparison revealed that NE cell density was higher in females compared to males (**Figure 1C**; t_10_=5.065, p=0.0005). In male subjects, in the entire LC, NE cell density was decreased in depressed suicides with a history of CA (DS-CA) compared to resilient individuals exposed to CA (**Figure 1D**; F_3,29_ = 4.060, p= 0.0159). Analysis of the effects of age, PMI, and pH on LC-NE cell density showed no significant correlations (**Supplementary Figure 2** and **3**).

**Table 1.** Group characteristics formalin (FRM) and frozen (FRZ) tissues. MDD: major depressive disorder; DD-NOS: depressive disorder not otherwise specified, PMI: post-mortem interval; SSRI: selective serotonin reuptake inhibitor; SNRI: selective norepinephrine reuptake inhibitor; TCA: tricyclic antidepressant.

| Formaline sample (FRM) |  |  |  |  |
| --- | --- | --- | --- | --- |
|  | CTL | DS | DS-CA | RES-CA |
| **N** | 10 | 10 | 12 | 9 |
| **Axis 1 diagnosis** | 0 | MDD (7)<br>DD-NOS (3) | MDD (8)<br>DD-NOS (4) | 0 |
| **Age (years)**<br>(*P=0.365*) | 58,27 +/- 5,12 | 46,90 +/- 3,89 | 42,58 +/- 4,02 | 48,78 +/- 4,00 |
| **Sex (M/F)** | 8/2 | 10/0 | 7/5 | 8/1 |
| **PMI (hours)**<br>(*P=0.0210*) | 51,13 +/- 3,53 | 68,39 +/- 6,23 | 71,99 +/- 6,64 | 50,73 +/- 6,87 |
| **pH tissue**<br>(*P=0.1379*) | 6,22 +/- 0,07 | 6,46 +/- 0,09 | 6,28 +/- 0,04 | 6,15 +/- 0,06 |
| **Substance at death** | TCA (1) ; Opioids (3) ;<br>SSRI (2) | TCA (1) ; SNRI (2) ;<br>Cocaine (1) ;<br>Canabinoids (2) ;<br>Alcohol (3) ;<br>Antipsychotic (1) | Canabinoids (2);<br>Cocaine (2);<br>Benzodiazepine (4);<br>Alcohol (3); SNRI (1);<br>TCA (1) | Canabinoids (2) ;<br>Opioids (3) ; Cocaine (2) ;<br>SSRI (2) ; Antipsychotic (2) |
| **Medication (last 3 months)** | Morphine (1) ;<br>Benzodiazepine (1) ;<br>SSRI (1) | Classic Antidepressant (2) ;<br>SNRI (1) ; Opioid (1) | SSRI (1); Sleeping Rx(1);<br>TCA (2); Benzodiazepine (1);<br>Antipsychotic (2);<br>Anticonvulsant (1);<br>Opioid (2) | Opioid analgesic(2) ;<br>Antidepressant (unknown 1) |
| Frozen sample (FRZ) |  |  |  |  |
|  | CTL | DS | DS-CA | RES-CA |
| **N** | 10 | 14 | 12 | 9 |
| **Axis 1 diagnosis** | 0 | MDD (10) DD-NOS (4) | MDD (10)<br>DD-NOS (2) | 0 |
| **Age (years)**<br>(*P=0.2281*) | 52,10 +/- 5,71 | 47, 29 +/- -3,69 | 41,54 +/- 2,91 | 51,00 +/- 4,12 |
| **Sex (M/F)** | 10/0 | 14/0 | 12/0 | 9/0 |
| **PMI (hours)**<br>(*P=0.3357*) | 58,65 +/- 5,32 | 69,49 +/- 5,83 | 73,58 +/- 6,84 | 62,19 +/- 6,08 |
| **pH tissue**<br>(*P=0.0679*) | 6,30 +/- 0,08 | 6,46 +/- 0,09 | 6,36 +/- 0,04 | 6,17 +/- 0,06 |
| **Substance at death** | TCA(1) ; Opioids (2) ;<br>SSRI (1) ; Amphetamine (1) | TCA (1) ; SNRI (2) ;<br>Cocaine (1) ;<br>Canabinoids (2) ;<br>Alcohol (3) ;<br>Antipsychotic (1) | Canabinoids (2);<br>Cocaine (2);<br>Benzodiazepine(4);<br>Alcohol (3); SNRI (1);<br>TCA (1) | Canabinoids(3) ;<br>Opioids (2) ; Cocaine (2) ;<br>SSRI (1)<br>Antipsychotic (2) |
| **Medication (last 3 months)** | Morphine (1) ;<br>Benzodiazepine (1) ;<br>SSRI (1) | Classic Antidepressant (2) ;<br>SNRI (1) ; Opioid (1) | SSRI (1); Sleeping Rx(1);<br>TCA (2);<br>Benzodiazepine (1);<br>Antipsychotic (2);<br>Anticonvulsant (1);<br>Opioid (2) | Opioid analgesic (2) ;<br>Antidepressant (unknown 1) |

We thus investigated whether specific regions of the LC (rostral vs. caudal) contributed to the observed decreased NE density in the DS-CA group (**Figure 1E**). We found that NE cell density was significantly decreased in the caudal part of the LC, but not in the rostral part, specifically in DS-CA (F_3,27_ = 5.44, p = 0.0047).

### Transcriptomic identity of LC-NE neurons in resilience

Our anatomical data suggest that protective mechanisms to the neuronal loss are taking place in resilient individuals to counteract the negative long-term effect of CA. To elucidate the molecular mechanisms underlying resilience, we performed a detailed transcriptomic analysis of LC-NE neurons. We employed LCM to isolate LC-NE neurons from frozen tissue samples (**Figure 2A**), followed by RNA sequencing to profile gene expression.

**Figure 2.**
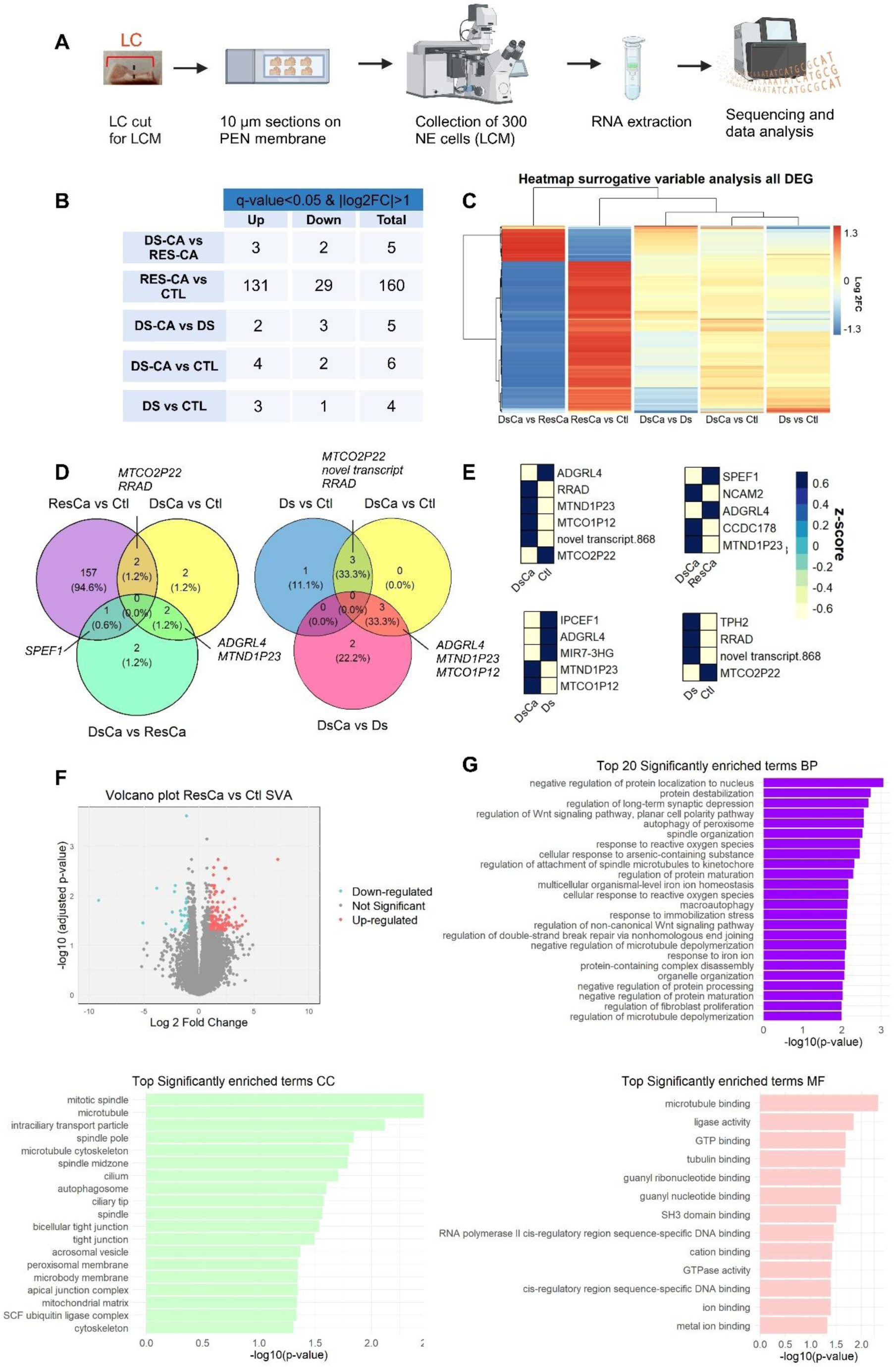
Specific profile of differentially expressed genes in child abuse, depression and resilience. **A.** Representative scheme of the experimental timeline: (1) the frozen block of locus coeruleus (LC) was cut in the middle to assure homogeneity between samples, (2) 5 to 6 sections of 10µm were collected and (3) 300 LC-NE cells per subject were collected on the LCM, (4) RNA extraction followed by (5) RNA sequencing and data analysis. **B.** Summary of the number of upregulated (Up) and downregulated (Down) differentially expressed genes (DEGs) for each comparison (RES-CA *vs* CTL, DS-CA *vs* CTL, DS-CA *vs* RES-CA, DS *vs* CTL, DS-CA *vs* DS). **C.** Heatmap of all DEGs illustrating global differences for all the comparisons. Data are presented as z-scores of Log2FC (scaled for each gene with a mean of 0 and a standard deviation of 1). **D.** Venn diagram comparing significantly differentially expressed genes between different conditions depicting gene overlapping. **E**. Heatmaps of DEGs between different groups with z-scores showing up-regulated genes in blue and down-regulated in yellow: number of DEG between DS-CA *vs* CTL – 6, DS-CA *vs* RES-CA – 5, DS-CA *vs* DS – 5, DS *vs* CTL – 4. **F.** Volcano plot of the comparison RES-CA *vs* CTL, with the most important number of DEGs. Up-regulated genes are depicted in orange, down-regulated genes in blue and the non-significant genes in gray. **G**. Summary of significantly enriched terms following topGO analysis in RES-CA *vs* CTL comparison of three ontologies: Biological Process (BP), Cellular Components (CC) and Molecular function (MF). In BP ontology, the top 20 significantly enriched terms were showed (only 4 more are illustrated in **Supplementary Figure 5**, **Supplementary Table 1**, due to the important number of significant enriched terms (118)). For CC and MF, the 19 and 13 significant terms, respectively, are presented.

The differential gene expression (DEG) patterns were assessed across the following comparisons: DS-CA vs CTL, RES-CA vs CTL, DS-CA vs RES-CA, DS vs CTL, and DS-CA vs DS. **Figure 2B** shows the number of upregulated and downregulated DEGs under stringent filtering criteria (|logFC≥1| and adjusted p-value ≤ 0.05). The RES-CA vs CTL comparison yielded the highest number of DEGs, totalling 160 (131 Upregulated and 29 Downregulated). The heatmap (**Figure 2C**) illustrates that the greatest changes in DEG fold change were observed in the RES-CA group relative to the CTL. An inverse relationship was noted between comparisons: genes upregulated in RES-CA compared to CTL were often downregulated in DS-CA compared to RES-CA, and vice versa (**Figure 2C**, **Figure 3**). For the remaining comparisons, our analysis identified a limited number of DEGs (3 genes common to DS individuals, 3 genes specific to DS-CA individuals and only one specific to the RES-CA individuals) (**Figure 2D**).

**Figure 3.**
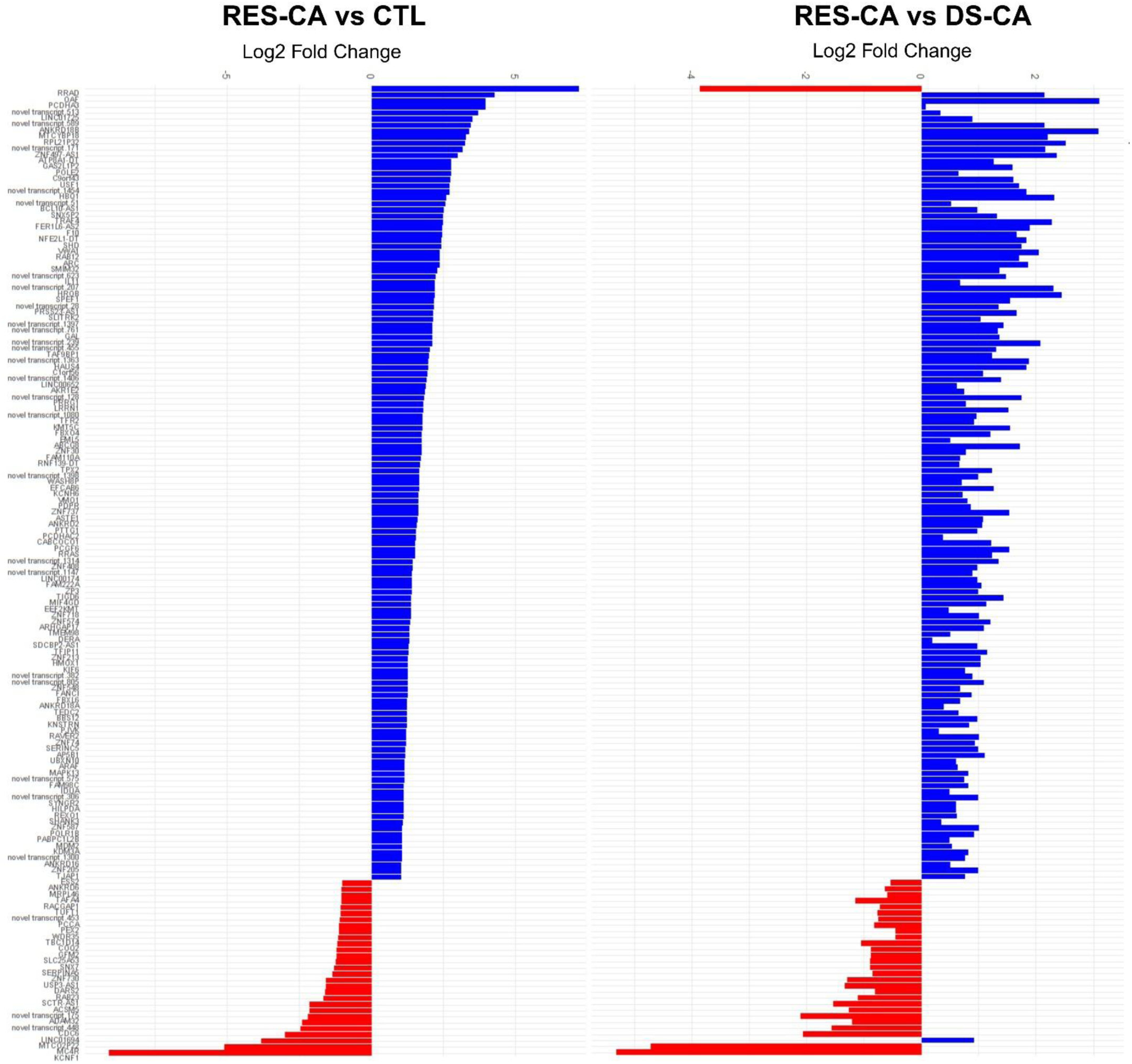
Histogram of the Log2 fold changes between RES-CA and CTL and between RES-CA and DS-CA, associated with the 160 differentially expressed genes (DEG) highlighted with adjusted p. value > 0.05 and |Log2FC| >1 in RES-CA *vs* CTL. Up-regulated genes are presented in blue and down-regulated genes in red.

Given the substantial number of DEGs identified in the RES-CA *vs* CTL comparison (**Figure 2E-F**, **Figure 3**), we conducted gene set enrichment analyses, only in this comparison. Significant findings were obtained from PsyGeNET (44), Pathfinder (45), and topGO. PsyGeNET, identified 13 out of 160 candidate genes involved in psychiatric disorders, highlighting roles in transport, receptor, oxidoreductase, and unclassified functions associated with schizophrenia, depression, bipolar disorder, and substance addictions (**data not shown**).

Pathfinder, revealed 26 significantly enriched pathways, which were portioned into 13 coherent clusters. Key pathways, with genes up-regulated in RES-CA, included (**Supplementary Figure 4A-B**): Ubiquitin mediated proteolysis (FBXO4 and MDM2), MAPK signaling pathway (RRAS, MAPK13, ARAF), Cell cycle (MDM2, PTTG1), FoxO signaling pathway (ARAF, MDM2, MAPK13) and Cellular senescence (MDM2, RRAS, MAPK13).

Finally, the topGO analysis identified significantly enriched terms across biological process (BP), cellular component (CC) and molecular function (MF). We found 118 BP terms, 19 CC terms and 13 MF terms that were significantly enriched (**Figure 2G, Supplementary Table 1**). To identify candidate’s genes, we explored the distribution between groups of genes associated with the top three terms in each category (**Supplementary Figure 5**). For BP enrichment, the analysis identified DEGs involved in negative regulation of protein localization to nucleus (FBXO4, RAB23 and TMEM89), protein destabilization (FBXO4, MDM2, PEX2) and regulation of long synaptic depression (ARK and SHANK3). While RAB23 and PEX2 were down-regulated in the RES-CA group, the other candidate genes were up-regulated. The three pathways the most significantly enriched for CC were mitotic spindle (KNRSTN, RACGAP1, TPX2, CDC6and HAUS4), microtubule (SPEF1, KNRTSN,EML5, FAMIIOA HAUS4, KIF6,

RACGAP1, TPX2), and intraciliary transport particles (UBXN10 and WDR35). Only RACGAP1 and CDC6 were down-regulated in the RES-CA group. Finally, the three pathways the most significantly enriched for MF were microtubule binding (EML5, HAUS4, KIF6, KNSTRN, RACGAP1and SPEF1), ligase activity (ACSM5, DARS2, MDM2 and PCCA), and GTP binding (RRAD, RAB12 and ACSM5).

### Transcriptomic profile of selected resilience-associated genes

To further investigate mechanisms linked to resilience, we examined whether genes of interest previously i) shown as enriched in LC-NE cells (33) or ii) associated with depression, CA or resilience in different context are significantly differentially expressed between the groups.

The LC-NE system is enriched in neuropeptides with some previously associated with resilience against stress in human and rodent models (9,46). We thus explored how neuropeptides-associated genes, and their receptors, were expressed specifically in NE cells (**Figure 4A-B**). Galanin (GAL; D_1_= 9.23, p= 0.0024), cortico-releasing hormone (CRH; D_1_= 4.31, p= 0.038), and enkephalin (PENK; D_1_= 7.60, p= 0.0058) were significantly overexpressed in the RES-CA group when compared to the CTL and/or DS-CA groups (**Figure 4A**). CRH was also found under expressed in DS-CA when compared to the DS group. The receptors genes for melanocortin (MC4R; D_1_= 4.55, p= 0.033) and Somatostatin (SSTR2; D_1_= 6.58, p= 0.010) were both under-expressed in RES-CA when compared to CTL or CTL and DS-CA, respectively (**Figure 4B**).

**Figure 4.**
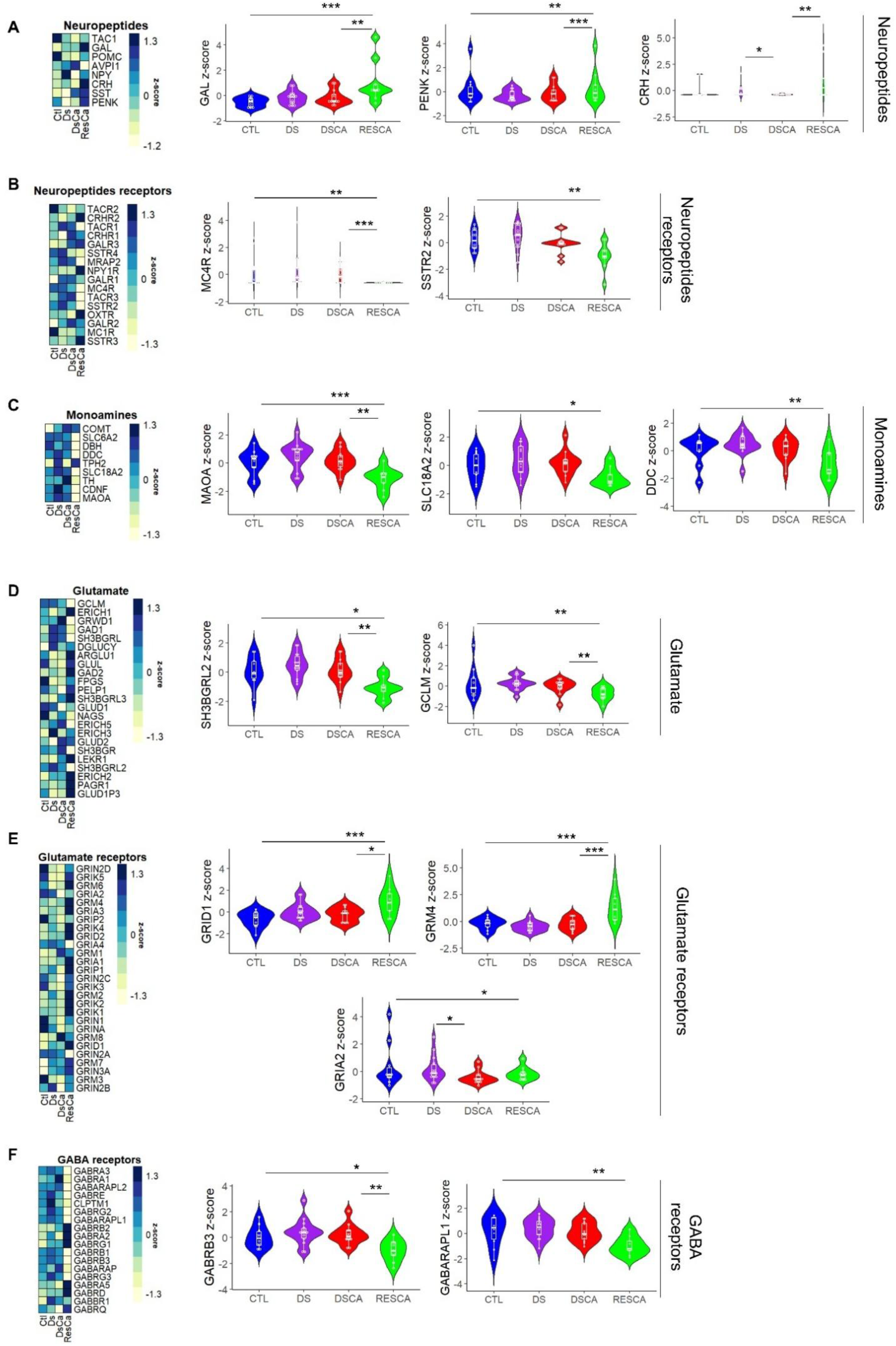
Genes expressed differentially in the noradrenergic cells of resilient individuals following child abuse. **A.** Heatmap of expression z-scores for selected neuropeptides in CTL, DS, DS-CA and RES-CA groups; Violin plots of normalized read counts for GAL (Galanin, D_1_= 9.23, p= 0.0024), CRH (cortico-releasing hormone, D_1_= 4.31, p= 0.038), and PENK (Enkephaline, D_1_= 7.6, p= 0.006). **B.** Heatmap of expression z-scores for selected neuropeptides receptors in CTL, DS, DS-CA and RES-CA groups; Violin plots of normalized read counts for MC4R (D_1_= 4.55, p= 0.033) and SSTR2 (D_1_= 6.58, p= 0.010). **C.** Heatmap of expression z-scores for monoamines in CTL, DS, DS-CA and RES-CA groups. Violin plot of normalized read counts for DDC (Dopa decarboxylase gene; D_1_= 6.71, p= 0.0096), SLC18A2 (vesicular monoamine transporter 2 gene; D_1_= 3.40, p= 0.046), MAOA (Monoamine Oxydase A gene; D_1_= 15.79, p= 0.000071) and HTR2A (5-hydroxytriptamine receptor 2A gene; D_1_= 5.61, p= 0.018). **D.** Heatmap of expression z-scores for Glutamate genes in CTL, DS, DS-CA and RES-CA groups. Violin plots of normalized read counts for GCLM (glutamate-cysteine ligase modifier subunit; D_1_= 7.88, p= 0.0050), SH3BGRL2 (SH3 Domain Binding Glutamate Rich Protein Like 2; D_1_= 7.17, p= 0.0074) and SH3BGRL3 (SH3 Domain Binding Glutamate Rich Protein Like 3; D(_1_)= 5.28, p = 0.022). **E.** Heatmap of expression z-scores for Glutamate receptors in CTL, DS, DS-CA and RES-CA groups. Violin plots of normalized read counts for GRIA2 (glutamate ionotropic receptor AMPA type subunit 2; D_1_= 6.29, p= 1.22e-02), GRM4 (glutamate metabotropic receptor 4; D(_1_)= 16.64, p= 4.53e-05) and GRID1 (glutamate ionotropic receptor delta type subunit 1; D_1_= 8.00, P = 4.66e-03). **F**. Heatmap of expression z-scores for GABA receptors in CTL, DS, DS-CA and RES-CA groups. Violin plots of normalized read counts of GABARAPL1 (GABA Type A Receptor Associated Protein Like 1; D_1_= 8.00, p= 0.0047) and GABRB3 (GABA Type A Receptor Subunit Beta3; D_1_= 6.40, p= 0.011). All the results are shown using boxplots showing 1st, 2nd (median) and the 3rd quartile (25%, 50% and 75%) of the distributions. When difference occurred according to the selected threshold, the difference between the groups is shown as followed: p-value (P) < 0.05 *, P< 0.01**, P< 0.001***.

Monoamines-associated genes involved in the synthesis, release and degradation of NE, serotonin (5HT) and dopamine (DA) and their receptors were studied. SLC18A2 (vesicular monoamine transporter 2 gene; D_1_= 3.40, p= 0.046), DDC (Dopa decarboxylase gene; D_1_= 6.71, p= 0.0096), MAOA (Monoamine Oxidase A gene; D_1_= 15.79, p= 0.000071) and HTR2A (5-hydroxytryptamine receptor 2A gene; D_1_= 5.61, p= 0.018) expression were down regulated in the RES-CA group, compared to CTL and/or DS-CA groups (**Figure 4C**). TPH2 (tryptophan hydroxylase 2 gene; D_1_= 15.43, p= 0.000086) was over-expressed in the DS group when compared to CTL and DS-CA (**data not shown**).

Since LC neurons are stimulated by glutamatergic inputs (47), we next focused on glutamate and glutamate receptor-associated genes (**Figure 4D-E**). Among these genes, ERICH1 (Glutamate-Rich Protein 1; D_1_= 5.17, p= 0.023) was specifically downregulated in depression without CA (**data not shown**). SH3BGRL3 (SH3 Domain Binding Glutamate Rich Protein Like 3; D_(1)_= 5.28, p = 0.022) and GRIA2 (glutamate ionotropic receptor AMPA type subunit 2; D_1_= 6.29, p= 1.22e-02) were respectively up- and down-regulated in CA. Regarding resilience, GCLM (glutamate-cysteine ligase modifier subunit; D_1_= 7.88, p= 0.0050) and SH3BGRL2 (SH3 Domain Binding Glutamate Rich Protein Like 2; D_1_= 7.17, p= 0.0074) were down regulated in RES-CA while GRM4 (glutamate metabotropic receptor 4; D(_1_)= 16.64, p= 4.53e-05) and GRID1 (glutamate ionotropic receptor delta type subunit 1; D_1_= 8.00, P = 4.66e-03) were up regulated in RES-CA compared to DS-CA and CTL (**Figure 4D-E**).

Although the LC is enriched in NE neurons, a group of GABAergic neurons in the dorsomedial area of the LC has been identified as a modulator of LC-NE neurons (48). We thus explore whether GABA signalling could be altered in LC-NE neurons, in the context of CA, depression and resilience (**Figure 4F**). Only two genes were down-regulated in RES-CA compared to CTL, GABARAPL1 (GABA Type A Receptor Associated Protein Like 1; D_1_= 8.00, p= 0.0047) and GABRB3 (GABA Type A Receptor Subunit Beta3; D_1_= 6.40, p= 0.011).

Finally, we compared genes from a recent spatially resolved transcriptomic study of LC neurons (**Supplementary Figure 6**). Most of the genes identified in Weber et al. (33) publication were present in our samples, except ARGAP36 and CCDC189. From this genes list, while overexpression of GDA (Guanine Deaminase; D_1_= 10.10, p= 0.0015) was associated with depression (without CA), it was under expressed in resilience. Moreover, CHODL (Chondrolectin; D_1_= 4.87, p= 0.027), LINC006682 (Long Intergenic Non-Protein Coding RNA 682; D_1_= 4.95, p= 0.026), RND3 (Rho Family GTPase 3; D_1_= 13.49, p= 0.00024) and GPX3 (Glutathione Peroxidase 3; D_1_= 4.91, p= 0.027) were under-expressed in RES-CA compared to CTL and/or DS-CA. Finally, PDGFD (Platelet Derived Growth Factor D; D_1_= 7.34, p= 0.0067) expression down-regulation was specifically associated with CA.

## Discussion

This study investigates the pathophysiology of depression, suicide and CA from an innovative and original angle, focusing on resilience. CA and early-life adversity are among the strongest lifetime predictors of depression and suicide. Examining resilience to MDD following CA is crucial for understanding protective and adaptive mechanisms preventing the emergence of long-lasting effects in the face of adversity, rather than merely identifying state markers. Increasing evidence implicates the LC-NE system in resilience following aversive experiences in both animals and humans at adulthood (28–31).

We aimed to explore, how a history of CA affects the integrity of the LC-NE system at molecular and cellular levels in human postmortem brain samples of depressed suicides, and whether differential neurobiological mechanisms can be identified in resilient individuals exposed to CA who did not develop any psychopathology. The anatomical study revealed decreased LC-NE density in depressed suicide individuals with a history of CA, an effect not seen in resilient individuals following CA (**Figure 1**). This suggests potential protective mechanisms in the LC of resilient individuals exposed to CA. To identify underlying molecular mechanisms, we performed RNA sequencing of LC-NE cells, revealing 160 genes with significant differential expression in the RES-CA group compared to controls (**Figure 2**), suggesting specific adaptations in LC-NE cells of resilient individuals. A summary of genes associated with LC-NE cells adaptation in resilience is illustrated **Figure 5**.

**Figure 5.**
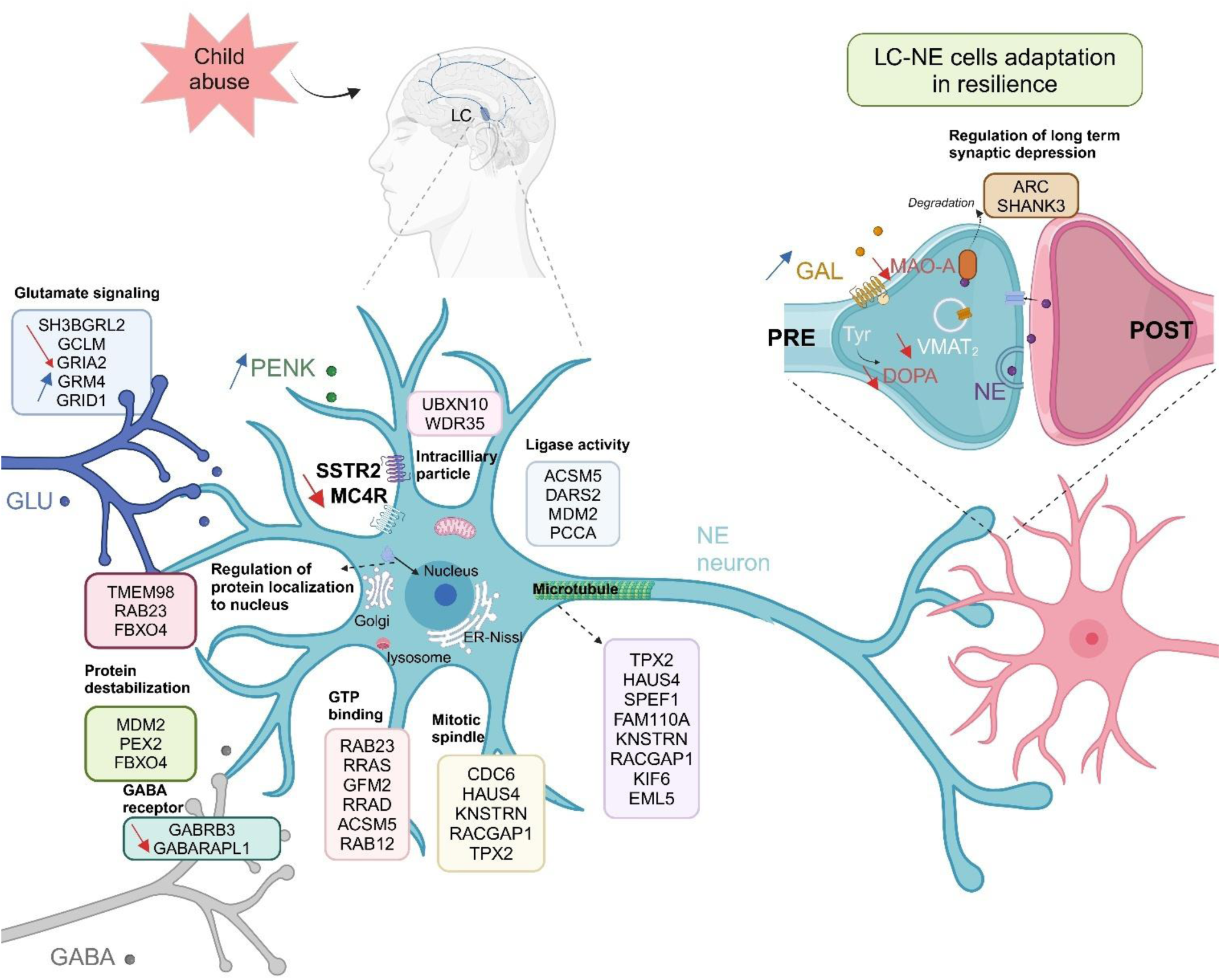
Summary image of the most significant pathways and differentially expressed genes between RES-CA and CTL. Created in Biorender. Sla, D. (2025), https://BioRender.com/o20c278; DOPA: Dihydroxyphenylalanine, Glu: Glutamate, GAL: Galanin, LC: Locus Coeruleus, MAO: Monoamine Oxydase, MC4R: Melanocortin 4 Receptor, NE: Noradrenaline, PENK: Proenkephalin, SSTR2: Somatostatin Receptor 2, Tyr: Tyrosine, VMAT2: Vesicular Monoamine Transporter 2.

Little is known about the impact of CA on the LC-NE system and how it affects the NE cells density. To our knowledge, this is the first study reporting anatomical changes in LC post-mortem tissues from depressed individuals with a history of CA. We found a significant decrease in LC-NE neuron density specifically in the caudal part of the LC in depressed suicide individuals with CA. Importantly, this effect was absent in depressed suicide without CA, highlighting that LC-NE loss is specific to CA-induced depression rather than depression from other etiological factors. When comparing this finding to the literature, studies examining TH expression in depressed suicide show inconsistent results: Arango et al. (49) found fewer neurons, Ordway et al. (50) reported increased TH levels, while Baumann et al. (51) found no change. Klimek et al. reported decrease NE transporter expression in depressive subjects in the mid-caudal LC but no difference in TH cells (52). Moreover, a magnetic resonance study reported intact LC integrity in late life depression (53). Differences in male and female subjects’ ratios across studies may explain these inconsistencies.

The LC is reported to be larger in female rats, with a greater number of NE neurons (54), and similar findings have been observed in humans (55). Our study showed comparable results when comparing male and female subjects, specifically within the CA-induced depression group (**Figure 1B**). However, due to the small female sample in other groups, sex-specific effects were excluded from our analyses.

Moreover, we found a specific decrease in NE cell density in the caudal LC of the DS-CA group. The caudal LC project to the spinal cord, cerebellum (via multipolar cells), hippocampus, and cortex (via fusiform cells) in rodents (56,57). Changes in hippocampal volume (6,58,59) and cortical areas, particularly the prefrontal cortex, have been associated not only to CA-induced depression (59–61), but also to resilience (15–18). This suggests the caudal LC-NE system may play a critical role in shaping long-term outcomes of CA, whether toward depression or resilience. A limitation of our study was the absence of rostral LC samples in some individuals (3 in DS, 1 in DS-CA, and 2 in RES-CA), preventing conclusions about NE cell density in that region.

Importantly, no LC-NE loss was observed in resilient individuals exposed to CA. It is crucial to note that resilience is not simply an absence of detrimental effects, but rather reflects active and adaptive processes protecting against long-term consequences of CA (9). This preservation of LC-NE density may be one mechanism that protect against the development of depression in adulthood.

Our transcriptomic profiling study represents the first post-mortem investigation exploring the molecular signatures of depression, CA, and resilience in human LC-NE cells. Only two previous studies investigated the molecular content of human LC-NE neurons, Barde et al. (2023) used LCM to study neuropeptide expression in control subjects (32) and Weber et al. (2024) provided insights into the spatial organization of the LC in control subjects (33).

In our study, we identified several genes associated with various conditions: overall depression (e.g., *RRAD*, *MTCO2P22*), CA-induced depression (e.g., *ADGRL4*, *MTCOP12*, *MTNDAP23*, *CRH*, *IPCEF1*, *MIR7-3HG*), depression without CA (e.g., *TPH2*, *ERICH1*, *GDA*), and genes associated specifically with CA (e.g., *SH3BGRL3*, *GRIA2*, *PDGFD*). These genes, not previously linked with depression or CA in large genetic study, may represent novel molecular markers specific to the NE system. The highest number of differentially expressed genes (DEGs) was found between the control and resilient groups, confirming significant adaptive changes in noradrenergic cells despite CA exposure.

Interestingly, only five DEGs distinguished the depressed suicide group with CA (DS-CA) from the RES-CA group, with *SPEF1* emerging as a resilience-specific gene. *SPEF1*, a microtubule-associated protein, play a key role in microtubule organisation, cell migration and projection organization. While no significant DEGs emerged between RES-CA and DS-CA among the genes highlighted between RES-CA and CTL, most genes upregulated in RES-CA compared to CTL were also upregulated in RES-CA compared to DS-CA, suggesting potential protective mechanisms (**Figure 3**).

Gene-set enrichment analysis revealed several significant gene categories in the RES-CA group. For biological processes, these included “negative regulation of protein localization to the nucleus,” “protein destabilization,” and “regulation of long-term depression.” cellular components like “mitotic spindle” and “microtubule” stood out, while molecular functions highlighted “ligase activity” and “GTP binding”. How the genes associated to these functions influence the integrity of LC-NE cells and the functional consequences of LC-NE signalling as potential markers of resilience remains to be investigated.

Among the selected genes (**Figure 4**), several showed specific dysregulation in resilient individuals, many previously associated with MDD in other brain regions or cell types. These include genes involved in monoamines metabolism and catabolism. *MAOA* responsible for monoamine degradation and *SLC18A2* (also known as VMAT2) linked to monoamine release showed dysregulation in resilient individuals. Surprisingly, *TPH2*, associated with serotonin synthesis, was dysregulated in NE neurons; its expression was specific to MDD in our dataset.

Additionally, neuropeptide-related genes were upregulated in resilient individuals following CA, particularly *Galanin* (*GAL*) and *pro-enkephalin* (*PENK*). *GAL,* expressed in approximately 80% of LC-NE cells (62), reduces LC firing rates when NE activity is elevated (63). Its increased expression in resilient individuals aligns with rodent models linking *GAL* to resilience, particularly after physical exercise (64,65). PENK, mainly found in A1 and A2 NE nuclei (66), has been detected in peri-LC GABAergic cells (67). Overexpression of δ-opioid receptors in the LC blocked stress-induced anxiety-like behaviour and LC firing properties (68).

We also observed changes in the regulation of NE neurons by other neurotransmitters, including serotonin (*HTR2A*), somatostatin (*SSTR2*) and melanocortin (*MC4R*), alongside GABA and glutamate. For instance, peri-LC GABAergic neurons modulate LC-NE neurons activity, but their role in depression and resilience remained unexplored, particularly in relation to anxiety-like behaviors or emotional regulation in rodent models.

One limitation of post-mortem studies is the inability to capture the dynamic nature of resilience, as they provide only a snapshot at the time of death. This makes it difficult to assess how neurobiological factors emerge or evolve over time, as well as their interactions. Additionally, drug effects on gene expression were challenging to evaluate. While antidepressants influence *GAL*, *MAOA*, and *TPH2* expression, only one subject per group was on antidepressants. Removing these individuals from the analysis did not alter results, suggesting observed gene expression changes were likely independent of medication. Similarly, cocaine and opioids, known to alter gene expression, had no apparent effects on identified genes.

In summary, while previous post-mortem studies have explored the long-term effects of CA using molecular and cellular approaches, our study uniquely identifies neurobiological markers specifically associated with resilience in LC-NE cells, rather than focusing solely on CA-induced changes. Beyond the anatomical reduction of NE cell density in depressed individuals with CA, we identified resilience-related biomarkers that could serve as potential therapeutic targets within noradrenergic cells, paving the way for future research.

## Acknowledgements

The authors express their gratitude to the financial support of the AAP AIN INSB CNRS (EI). The authors also acknowledge, for financial supports, the ANR – FRANCE (French National Research Agency, ANR-21-CE16-004, EI), the FRM (Fondation pour la recherche médicale, EQU202203014667, EI and BG), the Sissley Fundation, the IDEX Emergence grant (EI) and the doctoral school MCTI (ED 563, Médicament, Toxicologie, Chimie, Imageries) from the University Paris Cité, as well as the CIHR (Canadian Institute for Health Research, subvention N° 201909PJT-419517-PT, BG). The project was carried out with the support of the ERIE Foundation, a fund hosted by the King Baudouin Foundation (EI).

## Author contributions

D.S and E.I designed the experiments. D.S and M.A.D performed the anatomical study as well as laser capture microdissection and RNA extraction experiments. E.V performed brain slices at the cryostat for LCM. C.K performed the RNA sequencing analysis. G.F and C.N participated to the analysis. N.M and G.T furnished the human post-mortem samples from the Douglas brain bank. D.S and E.I wrote the manuscript that was edited by N.M, B.G, C.N and G.T. E.I supervised this research.

## Conflict of interest

The authors declare no conflict of interest.

**Supplementary information is available at MP’s website**

