## Supplementary information for "A post-mortem investigation of the locus coeruleus-noradrenergic system in resilience to childhood abuse"

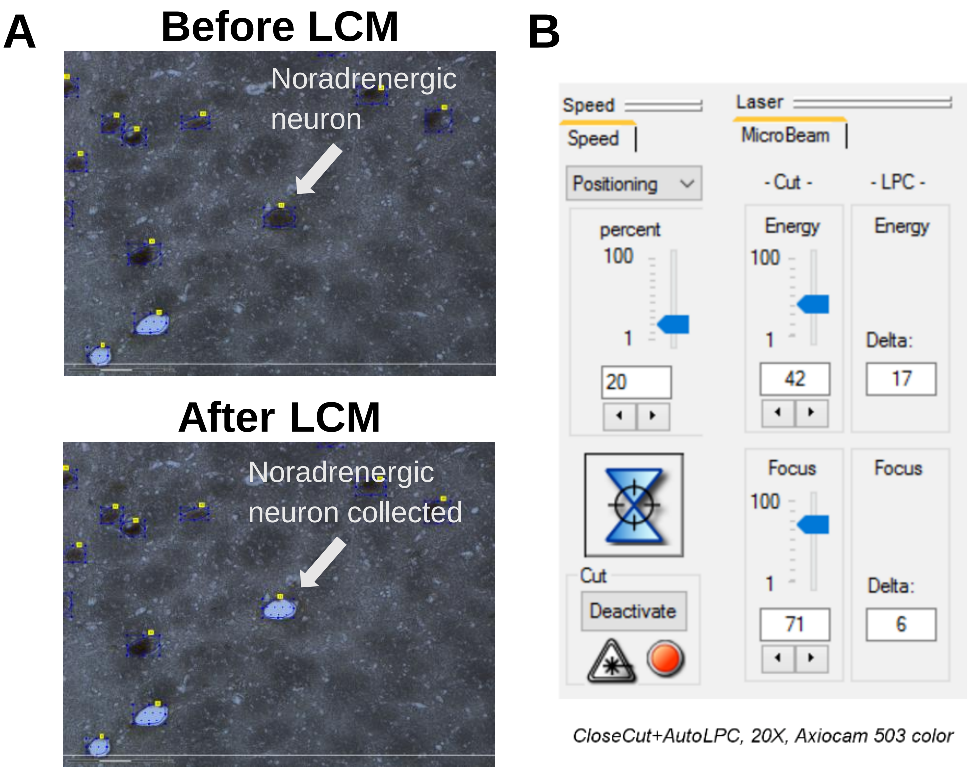


**Supplementary Figure 1. A**. Illustration of collection of noradrenergic (NE) neurons using laser capture microdissection (LCM). **Upper:** Representative image of NE neuron before being cut with the laser. **Lower:** The same neuron once the laser cut and collect the tissue. **B.** The parameters of the LCM used to cut and collect the 300 NE cells of each subject.

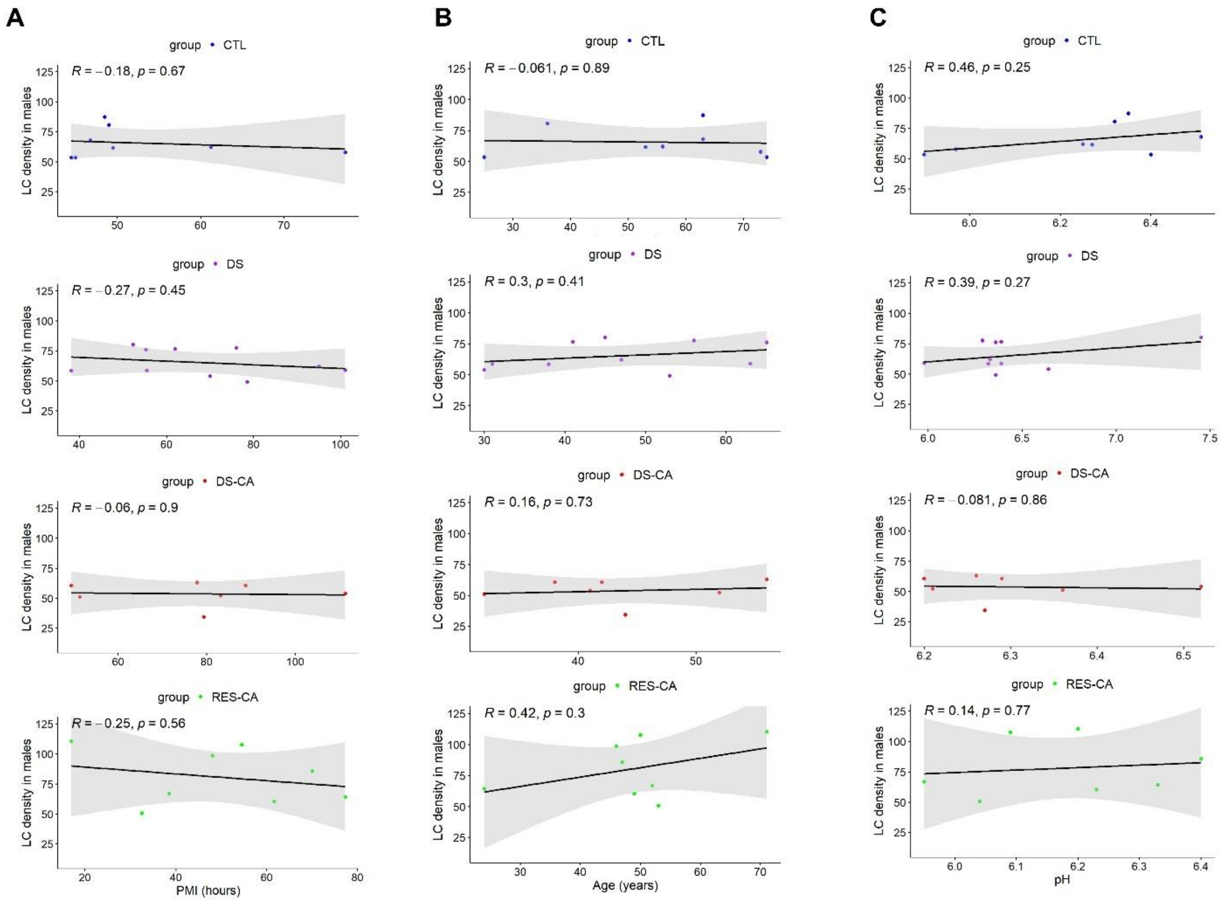


**Supplementary Figure 2. Correlative analysis of the effect of PMI, Age and pH on the density of whole LC in males. A.** Statistical analyses indicate no significant relationship between post-mortem interval (PMI) and TH+ cell density across all groups: Control (CTL) ($t_6$=-0.45, ρ= -0.18, p= 0.67) , Depression-Suicide (DS) ($t_8$=-0.79, ρ= -0.27, p= 0.457), Depression-Suicide-Child Abuse (DS-CA) ($t_5$=-0.137, ρ= -0.27, p= 0.45), and Child Abuse-No Depression (RES-CA) ($t_6$=-0.62, ρ= -0.25, p= 0.56) . **B**. No significant correlation is observed between age and the TH+ cells density across all groups: CTL ($t_6$=-0.15, ρ= -0.061, p= 0.89), DS ($t_8$=0.88, ρ= 0.30, p= 0.40), DS-CA ($t_5$=0.36, ρ= 0.16, p= 0.73), RES-CA ($t_6$=1.13, ρ= 0.42, p= 0.30). **C.** No significant correlation is observed between pH and the TH+ cells density across all groups: CTL ($t_6$=1.27, ρ= 0.46, p= 0.25), DS ($t_8$=1.19, ρ= 0.39, p= 0.27), DS-CA ($t_5$=-0.18, ρ= -0.081, p= 0.86), RES-CA ($t_6$=0.31, ρ= 0.14, p= 0.77). The graphs are presented for each condition (CTL in blue, DS in purple, DS-CA in red and RES-CA in green). In each graph, the coefficient of determination $R^2$ and p-value are shown.

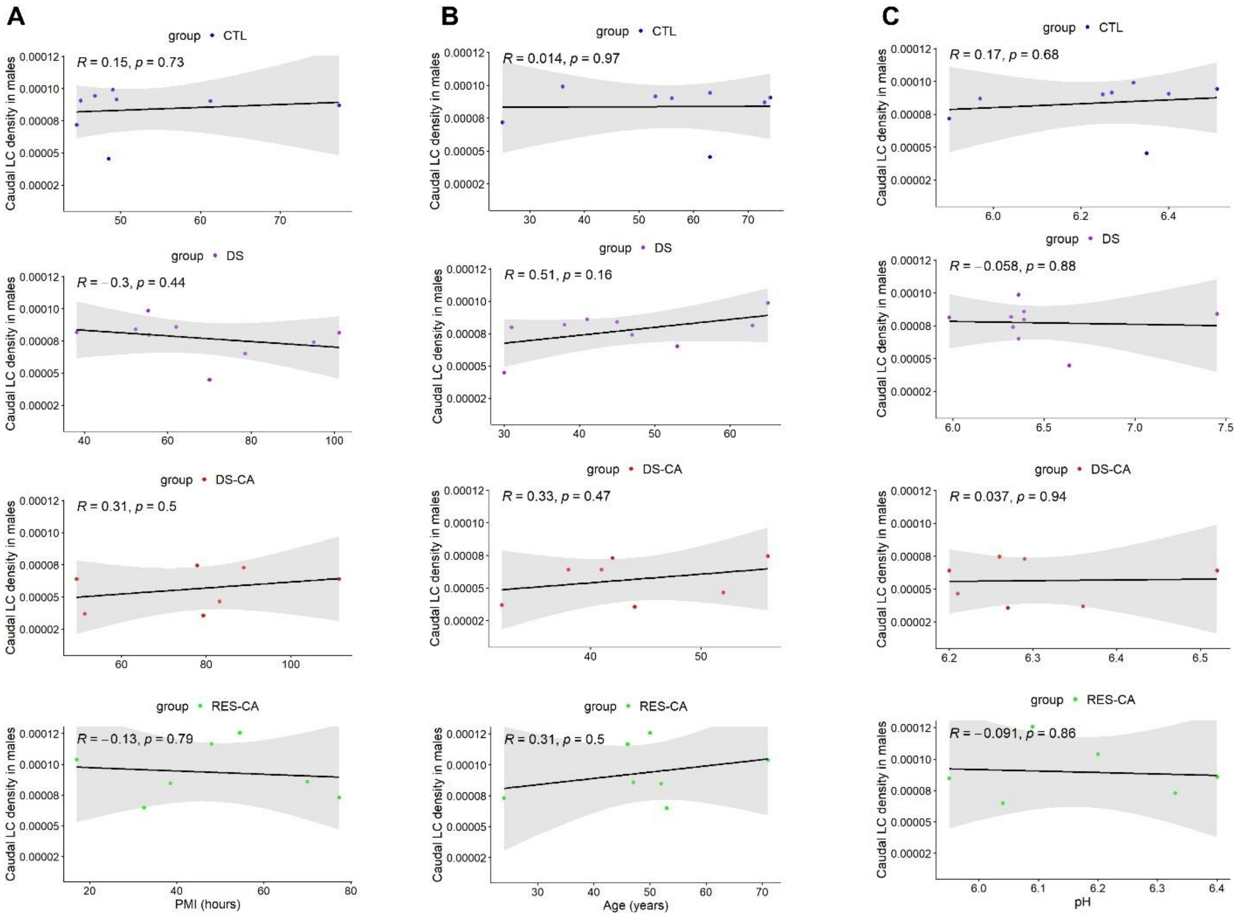

**Supplementary Figure 3. Correlative analysis of the effect of PMI, age and pH on the density of caudal LC in males. A**. Statistical analysis does not reveal significant correlation between the PMI and the caudal TH+ cells density across all groups: CTL ($t_6$= 0.36, ρ= 0.15, p= 0.73), DS ($t_7$= -0.83, ρ= -0.30, p= 0.44), DS-CA ($t_5$= 0.72, ρ= 0.31, p= 0.50), RES-CA ($t_4$= -0.28, ρ= -0.13, p= 0.79). **B**. No significant correlation between the age and the caudal TH+ cells density across all groups: CTL ($t_6$= 0.034, ρ= 0.014, p= 0.97), DS ($t_7$= 1.57, ρ= 0.51, p= 0.16), DS-CA ($t_5$= 0.79, ρ= 0.33, p= 0.47), RES-CA ($t_5$= 0.73, ρ= 0.31, p= 0.50). **C**. No significant correlation between the pH and the caudal TH+ cells density across all groups: CTL ($t_6$= 0.43, ρ= 0.17, p= 0.68), DS ($t_7$= -0.15, ρ= -0.058, p= 0.88), DS-CA ($t_5$= 0.084, ρ= 0.037, p= 0.94), RES-CA ($t_4$= -0.18, ρ= -0.091, p= 0.86). The graphs are presented for each condition (CTL in blue, DS in purple, DS-CA in red and RES-CA in green). In each graph, the coefficient of determination $R^2$ and p-value are shown.

**A**

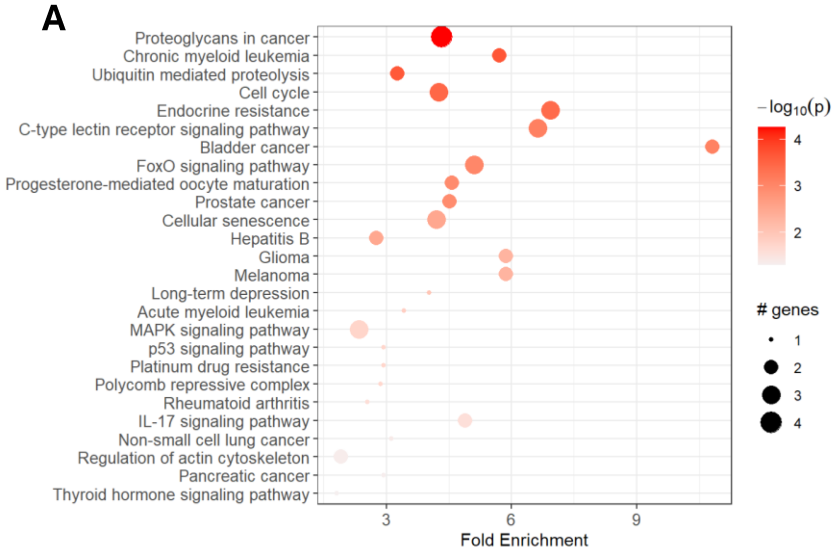


**B**

| ID | Term_Description | Fold_Enrichment | occurrence | support | lowest_p | highest_p | Up_regulated | Down_regulated |
|---|---|---|---|---|---|---|---|---|
| hsa05205 | Proteoglycans in cancer | 4.32956937799043 | 10 | 0.0306122448979592 | 5.49306180298756e-05 | 5.49306180298756e-05 | RRAS, ARAF, MDM2, MAPK13 | |
| hsa05220 | Chronic myeloid leukemia | 5.71262626262626 | 10 | 0.0547991924042344 | 0.000203862636150345 | 0.000203862636150345 | MDM2, ARAF | |
| hsa04120 | Ubiquitin mediated proteolysis | 3.26435786435786 | 10 | 0.0102040816326531 | 0.000223146489331307 | 0.00307342685475893 | FBXO4, MDM2 | |
| hsa04110 | Cell cycle | 4.25492163009404 | 10 | 0.0153061224489796 | 0.000340941237911513 | 0.000340941237911513 | MDM2, PTTG1 | CDC6 |
| hsa01522 | Endocrine resistance | 6.93217568947906 | 10 | 0.0737025851320274 | 0.000387529351094254 | 0.000387529351094254 | ARAF, MAPK13, MDM2 | |
| hsa04625 | C-type lectin receptor signaling pathway | 6.63401759530792 | 10 | 0.0153061224489796 | 0.000772311164854793 | 0.000772311164854793 | RRAS, MDM2, MAPK13 | |
| hsa05219 | Bladder cancer | 10.8239234449761 | 10 | 0.0837611118066859 | 0.000800800331687707 | 0.00546147353971666 | MDM2, ARAF | |
| hsa04068 | FoxO signaling pathway | 5.09887302779865 | 10 | 0.0255102040816327 | 0.000979579541013354 | 0.000979579541013354 | ARAF, MDM2, MAPK13 | |
| hsa04914 | Progesterone-mediated oocyte maturation | 4.57010101010101 | 10 | 0.0102040816326531 | 0.0011159241840296 | 0.0011159241840296 | MAPK13, ARAF | |
| hsa05215 | Prostate cancer | 4.51988011988012 | 10 | 0.0315868221994403 | 0.00115377533308803 | 0.00115377533308803 | MDM2, ARAF | |
| hsa04218 | Cellular senescence | 4.19703153988868 | 10 | 0.00510204081632653 | 0.00306477101979123 | 0.00306477101979123 | MDM2, RRAS, MAPK13 | |
| hsa05161 | Hepatitis B | 2.76046369737645 | 10 | 0.00510204081632653 | 0.00319160770885001 | 0.00319160770885001 | ARAF, MAPK13 | |
| hsa05214 | Glioma | 5.87584415584416 | 10 | 0.0500500813822461 | 0.00512505676285585 | 0.0187262729182985 | ARAF, MDM2 | |
| hsa05218 | Melanoma | 5.87584415584416 | 10 | 0.0500500813822461 | 0.00512505676285585 | 0.0187262729182985 | MDM2, ARAF | |
| hsa04730 | Long-term depression | 4.03244206773619 | 2 | 0.00502512562814070 | 0.00989763719360606 | 0.00989763719360606 | ARAF | |
| hsa05221 | Acute myeloid leukemia | 3.42757575757576 | 7 | 0.00510204081632653 | 0.0137329517729457 | 0.0137329517729457 | ARAF | |
| hsa04010 | MAPK signaling pathway | 2.33698347107438 | 10 | 0.00510204081632653 | 0.017637815077717 | 0.017637815077717 | RRAS, MAPK13, ARAF | |
| hsa04115 | p53 signaling pathway | 2.93792207792208 | 10 | 0.0102040816326531 | 0.0187262729182985 | 0.0187262729182985 | MDM2 | |
| hsa01524 | Platinum drug resistance | 2.93792207792208 | 10 | 0.00510204081632653 | 0.0187262729182985 | 0.0187262729182985 | MDM2 | |
| hsa03083 | Polycomb repressive complex | 2.85631313131313 | 10 | 0.0306122448979592 | 0.0198172675139542 | 0.0198172675139542 | PCGF6 | |
| hsa05323 | Rheumatoid arthritis | 2.53894500561167 | 10 | 0.00510204081632653 | 0.0251071311156646 | 0.0251071311156646 | IL11 | |
| hsa04657 | IL-17 signaling pathway | 4.8965367965368 | 10 | 0.00510204081632653 | 0.0270086267964271 | 0.0270086267964271 | MAPK13, TRAF4 | |
| hsa05223 | Non-small cell lung cancer | 3.11597796143251 | 2 | 0.00509067357512953 | 0.0414347779004608 | 0.0414347779004608 | ARAF | |
| hsa04810 | Regulation of actin cytoskeleton | 1.90420875420875 | 10 | 0.00510204081632653 | 0.0444432193445547 | 0.0444432193445547 | ARAF, RRAS | |
| hsa05212 | Pancreatic cancer | 2.93792207792208 | 2 | 0.00509067357512953 | 0.0466283233324967 | 0.0466283233324967 | ARAF | |
| hsa04919 | Thyroid hormone signaling pathway | 1.80398724082935 | 10 | 0.00510204081632653 | 0.0498148124137517 | 0.0498148124137517 | MDM2 | |

**Supplementary Figure 4. A.** Bubble chart of enrichment results of RES-CA *vs* CTL in the Pathfinder analysis (45) identified 26 enriched terms (there were only 133 out of 160 significantly different genes that were available in the database of Pathfinder). **B**. Table of the genes involved in the significantly enriched terms identified with Pathfinder; ID (e.g. hsa05205 refers to identify specific pathway in the data base that can be explored further).

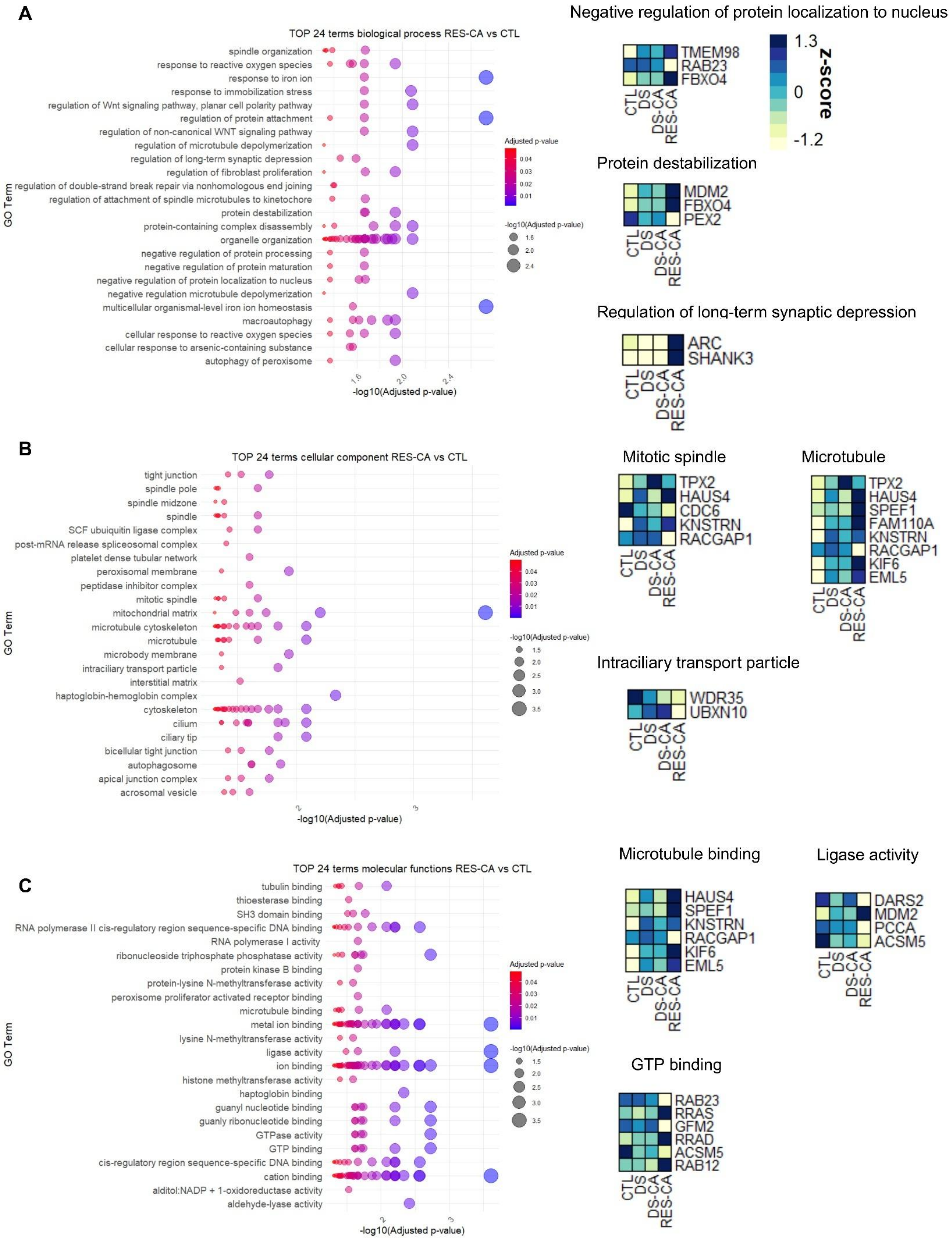

A
TOP 24 terms biological process RES-CA vs CTL
spindle organization
response to reactive oxygen species
response to iron ion
response to immobilization stress
regulation of Wnt signaling pathway, planar cell polarity pathway
regulation of protein attachment
regulation of non-canonical WNT signaling pathway
regulation of microtubule depolymerization
regulation of long-term synaptic depression
regulation of fibroblast proliferation
regulation of double-strand break repair via nonhomologous end joining
regulation of attachment of spindle microtubules to kinetochore
protein destabilization
protein-containing complex disassembly
organelle organization
negative regulation of protein processing
negative regulation of protein maturation
negative regulation of protein localization to nucleus
negative regulation microtubule depolymerization
multicellular organismal-level iron ion homeostasis
macroautophagy
cellular response to reactive oxygen species
cellular response to arsenic-containing substance
autophagy of peroxisome
GO Term
-log10(Adjusted p-value)
Adjusted p-value
-log10(Adjusted p-value)
Negative regulation of protein localization to nucleus
TMEM98
RAB23
FBXO4
CTL
DS
DS-CA
RES-CA
1.3
0
-1.2
z-score
Protein destabilization
MDM2
FBXO4
PEX2
Regulation of long-term synaptic depression
ARC
SHANK3
B
TOP 24 terms cellular component RES-CA vs CTL
tight junction
spindle pole
spindle midzone
spindle
SCF ubuiquitin ligase complex
post-mRNA release spliceosomal complex
platelet dense tubular network
peroxisomal membrane
peptidase inhibitor complex
mitotic spindle
mitochondrial matrix
microtubule cytoskeleton
microtubule
microbody membrane
intraciliary transport particle
interstitial matrix
haptoglobin-hemoglobin complex
cytoskeleton
cilium
ciliary tip
bicellular tight junction
autophagosome
apical junction complex
acrosomal vesicle
GO Term
-log10(Adjusted p-value)
Adjusted p-value
-log10(Adjusted p-value)
Mitotic spindle
TPX2
HAUS4
CDC6
KNSTRN
RACGAP1
Microtubule
TPX2
HAUS4
SPEF1
FAM110A
KNSTRN
RACGAP1
KIF6
EML5
Intraciliary transport particle
WDR35
UBXN10
C
TOP 24 terms molecular functions RES-CA vs CTL
tubulin binding
thioesterase binding
SH3 domain binding
RNA polymerase II cis-regulatory region sequence-specific DNA binding
RNA polymerase I activity
ribonucleoside triphosphate phosphatase activity
protein kinase B binding
protein-lysine N-methyltransferase activity
peroxisome proliferator activated receptor binding
microtubule binding
metal ion binding
lysine N-methyltransferase activity
ligase activity
ion binding
histone methyltransferase activity
haptoglobin binding
guanyl nucleotide binding
guanyl ribonucleotide binding
GTPase activity
GTP binding
cis-regulatory region sequence-specific DNA binding
cation binding
alditol:NADP + 1-oxidoreductase activity
aldehyde-lyase activity
GO Term
-log10(Adjusted p-value)
Adjusted p-value
-log10(Adjusted p-value)
Microtubule binding
HAUS4
SPEF1
KNSTRN
RACGAP1
KIF6
EML5
Ligase activity
DARS2
MDM2
PCCA
ACSM5
GTP binding
RAB23
RRAS
GFM2
RRAD
ACSM5
RAB12

**Supplementary Figure 5. A**. Bubble plot of genes in the top 24 biological process (BP) between RES-CA and CTL groups (**Left**). Heatmaps of expression z-scores for genes in the top 3 significantly expressed BP categories for the four different groups (CTL, DS, DS-CA and RES-CA) (**Right**). **B**. Bubble plot of genes in the top 24 cellular component (CC) between RES-CA and CTL groups (**Left**). Heatmaps of expression z-scores for genes in the top 3 significantly expressed CC categories for the four different groups (CTL, DS, DS-CA and RES-CA) (**Right**). **C**. Bubble plot of genes in the top 24 molecular functions (MF) between RES-CA and CTL groups (**Left**). Heatmaps of expression z-scores for genes in the top 3 significantly expressed MF categories for the four different groups (CTL, DS, DS-CA and RES-CA) (**Right**). The scale of z-score is the same for all heatmaps.

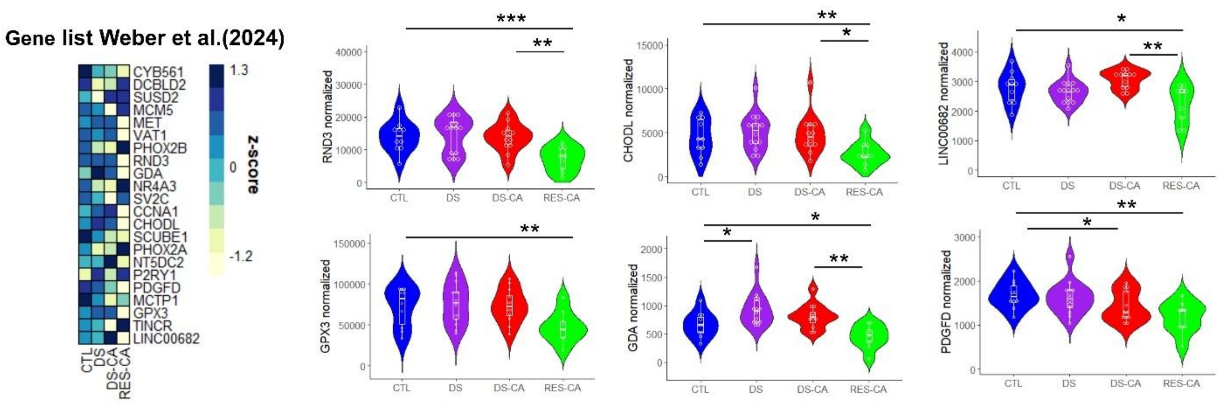


**Supplementary Figure 6.** Heatmap of expression z-scores for genes enriched in the LC-NE cells (Weber et al. 2024) in CTL, DS, DS-CA and RES-CA groups. Violin plots of normalized counts for RND3 (Rho Family GTPase 3; $D_1$= 13.49, p= 0.00024), GDA (Guanine Deaminase; $D_1$= 10.10, p= 0.0015), CHODL (Chondrolectin; $D_1$= 4.87, p= 0.027), PDGFD (Platelet Derived Growth Factor D; $D_1$= 7.34, p= 0.0067), GPX3 (Glutathione Peroxidase 3; $D_1$= 4.91, p= 0.027) and LINC00682 (Long Intergenic Non-Protein Coding RNA 682; $D_1$= 4.95, p= 0.026). All the results are shown using boxplots showing 1st, 2nd (median) and the 3rd quartile (25%, 50% and 75%) of the distributions. When difference occurred according to the selected threshold, the difference between the groups is shown as followed: p-value (P) < 0.05 *, P< 0.01**, P< 0.001***.

| Biological process (BP) | | | | | | |
|---|---|---|---|---|---|---|
| GO.ID | Term | Annotated | Significant | Expected | classicFisher | Significant_Genes |
| GO:1900181 | negative regulation of protein localization to nucleus | 38 | 3 | 0.19 | 0.00089 | FBXO4, RAB23, TMEM98 |
| GO:0031648 | protein destabilization | 49 | 3 | 0.24 | 0.00186 | FBXO4, MDM2, PEX2 |
| GO:1900452 | regulation of long-term synaptic depression | 14 | 2 | 0.07 | 0.00214 | ARC, SHANK3 |
| GO:0030242 | autophagy of peroxisome | 16 | 2 | 0.08 | 0.00281 | PEX2, PJVK |
| GO:2000095 | regulation of Wnt signaling pathway, planar cell polarity pathway | 16 | 2 | 0.08 | 0.00281 | ANKRD6, SPEF1 |
| GO:0007051 | spindle organization | 197 | 5 | 0.98 | 0.00304 | FAM110A, HAUS4, KNSTRN, RACGAP1, TPX2 |
| GO:0000302 | response to reactive oxygen species | 203 | 5 | 1.01 | 0.00346 | HMOX1, MAPK13, MDM2, PEX2, PJVK |
| GO:0071243 | cellular response to arsenic-containing substance | 18 | 2 | 0.09 | 0.00356 | HMOX1, MAPK13 |
| GO:0051988 | regulation of attachment of spindle microtubules to kinetochore | 21 | 2 | 0.1 | 0.00484 | KNSTRN, RACGAP1 |
| GO:1903317 | regulation of protein maturation | 70 | 3 | 0.35 | 0.00514 | MDM2, TFR2, TMEM98 |
| GO:0060586 | multicellular organismal-level iron ion homeostasis | 25 | 2 | 0.12 | 0.00682 | HMOX1, TFR2 |
| GO:0034614 | cellular response to reactive oxygen species | 151 | 4 | 0.75 | 0.00686 | MAPK13, MDM2, PEX2, PJVK |
| GO:0016236 | macroautophagy | 343 | 6 | 1.71 | 0.00734 | HMOX1, PEX2, PJVK, RAB23, SNX7, TBC1D14 |
| GO:0035902 | response to immobilization stress | 26 | 2 | 0.13 | 0.00737 | GAL, MDM2 |
| GO:0007026 | negative regulation of microtubule depolymerization | 27 | 2 | 0.13 | 0.00793 | SPEF1, TPX2 |
| GO:2000050 | regulation of non-canonical Wnt signaling pathway | 27 | 2 | 0.13 | 0.00793 | ANKRD6, SPEF1 |
| GO:2001032 | regulation of double-strand break repair via nonhomologous end joining | 27 | 2 | 0.13 | 0.00793 | KMT5C, TFIP11 |
| GO:0010039 | response to iron ion | 28 | 2 | 0.14 | 0.00852 | MDM2, TFR2 |
| GO:0032984 | protein-containing complex disassembly | 252 | 5 | 1.25 | 0.00852 | GFM2, PEX2, SPEF1, TFIP11, TPX2 |
| GO:0006996 | organelle organization | 3623 | 28 | 18.03 | 0.00872 | ARC, ARHGAP17, BBS12, CDC6, FAM110A, FBXO4, GFM2, HAUS4, KCNF1, KDM3A, KNSTRN, PEX2, PJVK, POLR1B, PTTG1, RAB12, RAB23, RACGAP1, SHANK3, SNX7, SPEF1, TBC1D14, TFIP11, TJAP1, TPX2, UBXN10, WDR35, ZNF205 |
| GO:0010955 | negative regulation of protein processing | 30 | 2 | 0.15 | 0.00974 | MDM2, TMEM98 |
| GO:1903318 | negative regulation of protein maturation | 30 | 2 | 0.15 | 0.00974 | MDM2, TMEM98 |
| GO:0048145 | regulation of fibroblast proliferation | 90 | 3 | 0.45 | 0.01026 | CDC6, FBXO4, PEX2 |
| GO:0031114 | regulation of microtubule depolymerization | 31 | 2 | 0.15 | 0.01037 | SPEF1, TPX2 |
| Cellular component (CC) | | | | | | |
| GO.ID | Term | Annotated | Significant | Expected | classicFisher | Significant_Genes |
| GO:0072686 | mitotic spindle | 184 | 5 | 0.97 | 0.0029 | CDC6, HAUS4, KNSTRN, RACGAP1, TPX2 |
| GO:0005874 | microtubule | 468 | 8 | 2.47 | 0.0034 | EML5, FAM110A, HAUS4, KIF6, KNSTRN, RACGAP1, SPEF1, TPX2 |
| GO:0030990 | intraciliary transport particle | 25 | 2 | 0.13 | 0.0076 | UBXN10, WDR35 |
| GO:0000922 | spindle pole | 177 | 4 | 0.93 | 0.0144 | CDC6, FAM110A, KNSTRN, TPX2 |
| GO:0015630 | microtubule cytoskeleton | 1405 | 14 | 7.42 | 0.0157 | CABCOCO1, CDC6, EML5, FAM110A, HAUS4, KIF6, KNSTRN, RAB23, RACGAP1, SERINC5, SPEF1, TEDC2, TPX2, WDR35 |
| GO:0051233 | spindle midzone | 37 | 2 | 0.2 | 0.0163 | CDC6, RACGAP1 |
| GO:0005929 | cilium | 764 | 9 | 4.03 | 0.0197 | BBS12, CABCOCO1, KCNF1, PJVK, SHANK3, SPEF1, TEDC2, UBXN10, WDR35 |
| GO:0005776 | autophagosome | 119 | 3 | 0.63 | 0.0252 | RAB12, RAB23, TBC1D14 |

| | | | | | | |
|---|---|---|---|---|---|---|
| GO:0097542 | ciliary tip | 48 | 2 | 0.25 | 0.0266 | SPEF1, WDR35 |
| GO:0005819 | spindle | 433 | 6 | 2.29 | 0.0273 | CDC6, FAM110A, HAUS4, KNSTRN, RACGAP1, TPX2 |
| GO:0005923 | bicellular tight junction | 126 | 3 | 0.67 | 0.0291 | ARHGAP17, TJAP1, TRAF4 |
| GO:0070160 | tight junction | 131 | 3 | 0.69 | 0.0322 | ARHGAP17, TJAP1, TRAF4 |
| GO:0001669 | acrosomal vesicle | 147 | 3 | 0.78 | 0.0429 | ARC, RACGAP1, SERPINA5 |
| GO:0005778 | peroxisomal membrane | 64 | 2 | 0.34 | 0.0450 | PEX2, PJVK |
| GO:0031903 | microbody membrane | 64 | 2 | 0.34 | 0.0450 | PEX2, PJVK |
| GO:0043296 | apical junction complex | 150 | 3 | 0.79 | 0.0451 | ARHGAP17, TJAP1, TRAF4 |
| GO:0005759 | mitochondrial matrix | 491 | 6 | 2.59 | 0.0458 | ACSM5, DARS2, GFM2, MRPL46, PCCA, PDPR |
| GO:0019005 | SCF ubiquitin ligase complex | 65 | 2 | 0.34 | 0.0463 | FBXL6, FBXO4 |
| GO:0005856 | cytoskeleton | 2424 | 19 | 12.79 | 0.0495 | ARC, CABCOCO1, CDC6, EFCAB6, EML5, FAM110A, HAUS4, KIF6, KNSTRN, PJVK, RAB23, RACGAP1, SERINC5, SPEF1, TEDC2, TPX2, TRAF4, WDR35, ZNF74 |
| **Molecular function (MF)** | | | | | | |
| **GO.ID** | **Term** | **Annotated** | **Significant** | **Expected** | **classicFisher** | **Significant_Genes** |
| GO:0008017 | microtubule binding | 277 | 6 | 1.56 | 0.0047 | EML5, HAUS4, KIF6, KNSTRN, RACGAP1, SPEF1 |
| GO:0016874 | ligase activity | 166 | 4 | 0.93 | 0.0143 | ACSM5, DARS2, MDM2, PCCA |
| GO:0005525 | GTP binding | 380 | 6 | 2.13 | 0.0203 | ACSM5, GFM2, RAB12, RAB23, RRAD, RRAS |
| GO:0015631 | tubulin binding | 383 | 6 | 2.15 | 0.0210 | EML5, HAUS4, KIF6, KNSTRN, RACGAP1, SPEF1 |
| GO:0019001 | guanyl nucleotide binding | 401 | 6 | 2.25 | 0.0256 | ACSM5, GFM2, RAB12, RAB23, RRAD, RRAS |
| GO:0032561 | guanyl ribonucleotide binding | 401 | 6 | 2.25 | 0.0256 | ACSM5, GFM2, RAB12, RAB23, RRAD, RRAS |
| GO:0017124 | SH3 domain binding | 122 | 3 | 0.69 | 0.0314 | ARHGAP17, PTTG1, SHANK3 |
| GO:0000978 | RNA polymerase II cis-regulatory region sequence-specific DNA binding | 1194 | 12 | 6.7 | 0.0354 | USF1, ZNF205, ZNF213, ZNF30, ZNF408, ZNF548, ZNF574, ZNF587, ZNF718, ZNF730, ZNF737, ZNF74 |
| GO:0043169 | cation binding | 4391 | 33 | 24.66 | 0.0379 | ABCG8, ACSM5, ARAF, CABCOCO1, EFCAB6, F10, HBQ1, HMOX1, KDM3A, KMT5C, MDM2, PCCA, PCDHA3, PCDHAC2, PCGF6, PEX2, POLR1B, PRRG1, RACGAP1, SERPINA5, SHANK3, TRAF4, ZNF205, ZNF213, ZNF30, ZNF408, ZNF548, ZNF574, ZNF587, ZNF718, ZNF730, ZNF737, ZNF74 |
| GO:0003924 | GTPase activity | 333 | 5 | 1.87 | 0.0397 | GFM2, RAB12, RAB23, RRAD, RRAS |
| GO:0000987 | cis-regulatory region sequence-specific DNA binding | 1218 | 12 | 6.84 | 0.0403 | USF1, ZNF205, ZNF213, ZNF30, ZNF408, ZNF548, ZNF574, ZNF587, ZNF718, ZNF730, ZNF737, ZNF74 |
| GO:0043167 | ion binding | 6070 | 43 | 34.08 | 0.0407 | ABCG8, ACSM5, ARAF, BBS12, CABCOCO1, CDC6, DARS2, EFCAB6, F10, GFM2, HBQ1, HMOX1, KDM3A, KIF6, KMT5C, MAPK13, MDM2, PCCA, PCDHA3, PCDHAC2, PCGF6, PEX2, POLR1B, PRRG1, RAB12, RAB23, RACGAP1, RRAD, RRAS, SERPINA5, SHANK3, TRAF4, ZNF205, ZNF213, ZNF30, ZNF408, ZNF548, ZNF574, ZNF587, ZNF718, ZNF730, ZNF737, ZNF74 |
| GO:0046872 | metal ion binding | 4300 | 32 | 24.14 | 0.0465 | ABCG8, ACSM5, ARAF, CABCOCO1, EFCAB6, F10, HBQ1, HMOX1, KDM3A, KMT5C, MDM2, PCCA, PCDHA3, PCDHAC2, PCGF6, PEX2, POLR1B, PRRG1, RACGAP1, SHANK3, TRAF4, ZNF205, ZNF213, ZNF30, ZNF408, ZNF548, ZNF574, ZNF587, ZNF718, ZNF730, ZNF737, ZNF74 |

**Supplementary Table 1**. Biological process (BP), Cellular component (CC) and Molecular function (MF) significantly enriched GO terms following topGO enrichment analysis. Due to the important number of enriched terms in the BP category (118), we showed only the first 24.
